# Heat dissipation from photosynthesis contributes to maize thermoregulation under suboptimal temperature conditions

**DOI:** 10.1101/2023.01.27.525868

**Authors:** Verónica Sobejano-Paz, Xingguo Mo, Suxia Liu, Teis Nørgaard Mikkelsen, Lihong He, Hongxiao Jin, Mónica García

**Affiliations:** Key Laboratory of Water Cycle and Related Land Surface Processes, Institute of Geographical Sciences and Natural Resources Research, Chinese Academy of Sciences, Beijing, China.; Department of Environmental and Resource Engineering, Technical University of Denmark, Kgs. Lyngby, Denmark; Sino-Danish College, University of Chinese Academy of Sciences, Beijing, China; Sino-Danish Center for Education and Research, Aarhus, Denmark; CEIGRAM (Center for Environmental and Agricultural Risks). ETSIAAB. Universidad Politécnica de Madrid, 28040, Madrid, Spain.; Department of Physical Geography and Ecosystem Science, Lund University, 22362, Lund Sweden

**Keywords:** thermoregulation, Non-Photochemical Quenching (NPQ), thermal remote sensing, water stress, fluorescence, stomatal conductance, photosynthesis, optimum temperature, heat waves, maize, transpiration

## Abstract

The extent to which plants thermoregulate to maintain relatively stable metabolic function in response to gradual and rapid temperature changes that jeopardize crop production is unclear. Maize thermoregulation was investigated based on leaf temperature (T_L_) measurements and its relationship with photochemistry and stomatal conductance (g_s_) under dry and wet soil scenarios. Seasonal climatology was simulated in a growth chamber according to Beijing’s climatology with extreme “hot days” based on historical maxima.

Maize behaved as a limited homeotherm, an adaptive strategy to maintain photosynthesis around optimum temperatures (T_opt_). Plants on drier soil had lower thermoregulatory capacity, with reduced g_s_, photosynthesis and transpiration, which impacted final yields, despite acclimation with a higher T_opt_ to sustained stress. On hot days thermoregulation was affected by heat stress and water availability, suggesting that strong and frequent heatwaves will reduce crop activity although increased temperatures could bring photosynthesis closer to T_opt_ in the region.

We propose a novel mechanism to explain thermoregulation from the contribution of heat dissipation via non-photochemical quenching (NPQ) to T_L_, supporting our hypothesis that NPQ acts as a negative feedback mechanism from photosynthesis by increasing T_L_ in suboptimal conditions. These results could help to design adaptation strategies based on deficit irrigation.

**Highlight:** Maize was able to maintain leaf temperatures in narrower ranges than air temperatures by dissipating sunlight not used in photosynthesis as heat energy with a key role of transpiration cooling to sustain optimum photosynthesis temperature.

## 1. Introduction

Heatwaves and droughts are increasing in frequency, intensity and duration (IPCC, 2020) hampering crop production (Seneviratne et al., 2012; Teuling, 2018). Maize as one of the most important cereals for global food production, covering about 139 million ha (FAO, 2020), is already experiencing crop losses attributed to water and heat stress (Aslam et al., 2015; Liu et al., 2010; Lobell et al., 2013; Mishra and Cherkauer, 2010). Heatwaves are often followed by soil drought, which amplifies physiological damage (Das et al., 2016; Luan and Vico, 2020) complicating the understanding of plant responses to individual stressors. In fact, plants deploy different adaptive strategies to avoid or tolerate heat (Lobell et al., 2013; Tiwari and Yadav, 2019) or water stress (Gerhards et al., 2016; Leinonen and Jones, 2004; Sobejano-Paz et al., 2020). Disentangling maize responses to heatwaves and droughts is crucial to design adaptation strategies such as cooling irrigation or cultivar selection (Brás et al., 2021; Li et al., 2020).

Currently, it is unclear to what extent plants thermoregulate, buffering steady and sudden increases in air temperature (T_air_) to maintain relatively stable metabolic function (Geange et al., 2020). It is hypothesized that leaf traits have evolved to thermoregulate, maintaining leaf temperature (T_L_) within specific ranges to optimize carbon gains at instantaneous and lifetime scales (Fauset et al., 2018; Michaletz et al., 2016, 2015) as enzymatic reactions and membrane processes depend on T_L_ (Lambers et al., 2008; Schulze et al., 2005). To what extent thermoregulation occurs is not clear, with most of studies focusing in forests (Still et al., 2022; Drake et al., 2020; Fauset et al., 2018).

Based on the relationship between T_L_ and T_air_, plants could be classified as homeotherms (constant T_L_) when the T_air_ vs T_L_ slope (referred to as *β*) is 0, or poikilotherms with T_L_ mirroring variations in T_air_ (*β* close to 1). Previous research has confirmed that several plants tend to behave as limited homeotherms with *β* around 0.7 (Michaletz et al., 2016). If there is thermoregulation, the temperature at which the leaf temperature excess (ΔT = T_L_ - T_air_) is zero, called the leaf-air equivalence temperature (T_eq_), should become close to an optimum temperature (T_opt_) at which photosynthesis rate reaches maximum (Drake et al., 2020; Michaletz et al., 2016) but to our knowledge, thermoregulation and its mechanisms have not been specifically studied for maize.

It is well established that when T_L_ is below the T_opt_, photosynthesis tends to increase with T_L_ (Moore et al., 2021; Yamori et al., 2014). When T_L_ is above the T_opt_, photosynthesis is inhibited due to enzyme degradation (Guha et al., 2018; Killi et al., 2017; Moore et al., 2021). The T_opt_ has been identified for maize between 33 and 38°C (Crafts-Brandner and Salvucci, 2002; Hatfield et al., 2011; Moore et al., 2021). However, the T_opt_ varies with crop type, genotype, phenological stage or environmental conditions (Schulze et al., 2019; Yamori et al., 2014). In fact, plant acclimation under heat or water stress helps to maintain or enhance photosynthesis by increasing T_opt_ (Lambers et al., 2008; Vico et al., 2019; Zheng et al., 2018), contributing to heat tolerance (Tiwari and Yadav, 2019).

T_L_ can be used to evaluate plant thermoregulation and acclimation, presenting great potential for remote sensing thermal applications (Farella et al., 2022). T_L_ depends on several interrelated factors. In the short term (e.g. minutes to days), T_L_ results from balancing leaf energy budget components: from absorbing and scattering radiation in the shortwave range, absorbing and emitting longwave radiation and from dissipation of heat as sensible (H) or latent heat (or evapotranspiration) fluxes (λE) (Bonan, 2015; Jones and Rotenberg, 2001). In general, plants are more capable to reduce their temperature, mostly via transpiration, than to increase it when it is below T_opt_ (Mahan and Upchurch, 1989).

A direct but unexplored mechanism connecting leaf thermoregulation and photosynthesis is the increase in regulatory thermal dissipation as a photoprotective mechanism of photosystem II due to losses in the electron transport system. From the absorbed photosynthetic active radiation (PAR) around 20-50% can be used for photosynthesis depending on environmental and plant conditions. To avoid damage of photosystems, the rest of absorbed energy has to be dissipated. While less than 5% of the energy is reemitted as fluorescence, the largest part, around 80-50%, is dissipated as heat, which can be evaluated as non-photochemical quenching (NPQ) (Demmig-Adams et al., 2006; Endo et al., 2014). NPQ as an indicator of heat dissipation includes constitutive thermal dissipation and variable energy-dependent heat dissipation (Chen et al., 2019; Van der Tol et al., 2014) but the actual contribution of NPQ to T_L_ is not known. When T_L_ is below T_opt_, the proportion of absorbed photons dissipated as NPQ increased for rice cultivars (Hirotsu et al., 2004; Tsonev and Hikosaka, 2003). Kaňa and Vass (2008) showed experimentally that T_L_ increased proportionally to NPQ under constant T_air_, as photosynthesis decreased in model plant *Arabidopsis*.

Even though various photochemical parameters like the quantum yield of photosynthesis (Φ_P(II)_), the maximum photosynthetic quantum efficiency in the dark (F_v_/F_m_) or the Sun Induced Fluorescence (SIF) have been used to quantify plant responses to heat (Geange et al., 2020; Martini et al., 2022; Murchie and Lawson, 2013). To our knowledge, the actual contribution of NPQ to leaf thermoregulation via energy budgets has not been investigated. Its estimates can be used to assess photosynthetic efficiency using thermal remote sensing data with subsequent implications for predicting temperature-dependent processes such as carbon and water fluxes or leaf mortality (Blonder and Mitcheletz, 2018).

Another key variable regulating T_L_ especially through transpiration cooling is stomatal conductance (g_s_). Stomata close in response to hydraulic signals of soil and atmospheric drought resulting in higher T_L_ (Grossiord et al., 2020; Jones and Rotenberg, 2001). There are two opposite effects of higher vapor pressure deficit (VPD) on leaf transpiration caused by a warmer atmosphere. On one hand, the higher vapor diffusivity increases transpiration rates (T_r_) (Crafts-Brandner and Salvucci, 2002; Duursma et al., 2014; Guha et al., 2018; Tan et al., 2017; Zheng et al., 2018). On the other hand, stomata counteract this effect by closing and decreasing T_r_ when VPD reaches a certain threshold. Stomata also respond directly to changes in photosynthesis (Drake et al., 2020; Duursma et al., 2014; Moore et al., 2021) and possibly to changes in T_air_ (Urban et al., 2017) showing contradictory responses to increasing T_L_ in various studies (Moore et al., 2020). This means that assessing thermoregulation based on current g_s_ models remains challenging, especially when T_opt_ is unknown (Still et al,. 2022).

This study aims to explore leaf thermal dynamics and its association with photochemical responses, and to reveal adaptive strategies of T_L_ to extreme hot days (with moderate water stress and without water stress) using manipulative experiments on maize. We tested the hypothesis that heat dissipation from NPQ acts as a negative feedback mechanism from photosynthesis increasing warming towards T_opt_ in suboptimal conditions.

We focused on the following particular objectives:

1. To what extent do maize plants thermoregulate under wet and dry soil conditions during a climatic average growing season?
2. What is the role of photochemistry and stomatal conductance in thermoregulation during a climatic average growing season?
3. How does thermoregulation during hot days (temperature extremes) differ from that happening over normal or climatic average days?

We conducted a maize experiment in near-field conditions, where average changes in T_air_ were imposed to plants. Extreme events were imposed as a sudden T_air_ shocks based on the absolute maximum historical T_air_ records in each plant growth stage at the region.

## 2. Materials and methods

### 2.1. Experimental Design

The experiment was carried out from October 2018 to February 2019 using the Water Transformation Dynamical Processes Experimental Device (WATDPED) at the Institute of Geographic Sciences and Natural Resources Research (IGSNRR), Chinese Academy of Sciences, in Beijing. The WATDPED is a growth chamber containing two lysimeters with a ground area of 6 m^2^ and a depth of 3 m, reaching a capacity of 18 m^3^ each (Figure 1). The substrate is a mixture of sand, silt, and clay, homogenously distributed in one of the lysimeters and stratified in the other. The surface soil field capacity (SFC) was about 0.3 cm^3^/cm^3^, measured before crop sowing. The total weight of each lysimeter was recorded every 30 minutes using a datalogger. A 3×4 light matrix (6 metal halide and 6 sodium lamps) of 30 Klux illuminance capacity illuminates each lysimeter uniformly over the soil area and provides a downwelling quantum irradiance of about 800-1000 μmol m^−2^ s^−1^, similar to Jimenez et al. (2012). Distance between the lamps and ground was adjusted with a pulley system, keeping the lamps 2m above the top of the canopy to avoid photodamage. Environmental conditions such as T_air_, relative humidity (RH), CO_2_ concentration and light intensity were recorded every 3 minutes. Ventilation systems and centrifugal humidifiers are placed along two sides inside the growth chamber to control T_air_ and RH (Figure 1B). The successful use of the WATDPED has been reported in previous maize physiology studies (Chen et al., 2019; Tan et al., 2017; Wu et al., 2017).

**Figure 1.**
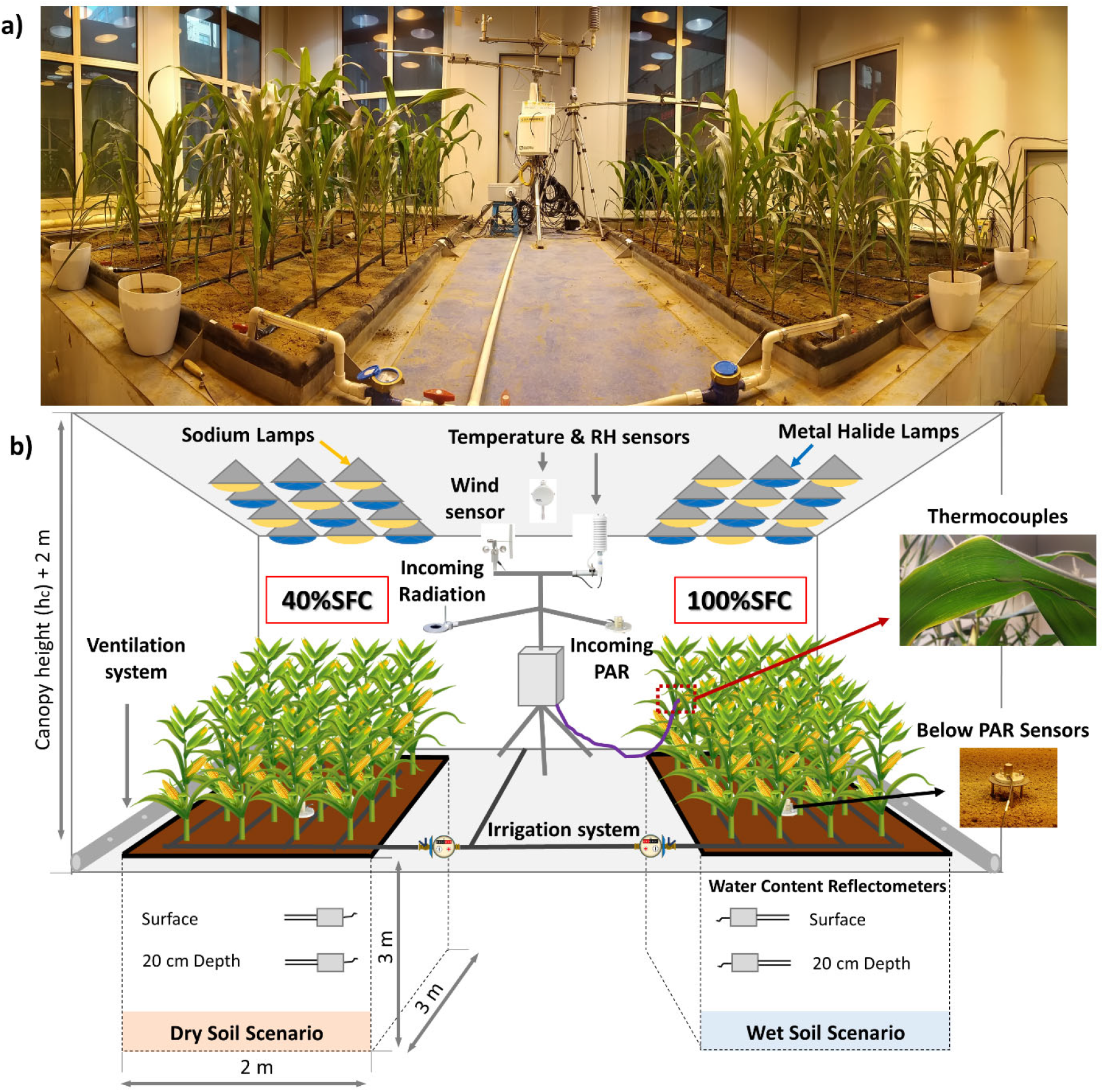
a) Panoramic view of the Water Transformation Dynamical Processes Experimental Device (WATDPED) after one month sowing. b) Schematic diagram of the WATDPED and main components. In the ground level, light racks of sodium (clear yellow) and metal halide lamps (dark blue) are uniformly distributed. The two lysimeters are named by their field capacity at the surface layer (SFC): Wet Soil Scenario (100%SFC) and Dry Soil Scenario (40%SFC). The irrigation system is manually controlled, and indoor climate is regulated through side ventilation.

Seeds of summer hybrid maize (*Zea mays* L. Zhengdan 958) were sown on the 12^th^ of October 2018, at approximate 5 cm soil depth with an inner space of 37 cm and 54 cm in the horizontal plane, allocating 36 plants in each lysimeter (Figure 1). To promote rapid growth and simulate daily illumination conditions during the growing season in Beijing, the light source was kept on between 6:00 h. and 19:00 h., simulating 13 hours of daytime. T_air_ and RH were set to simulate the climatology of Beijing during maize growing season (from 15^th^ of June to 15^th^ of October between 2000 and 2016). T_air_ was elevated on eight specific dates (referred to as “hot days”) for about two hours during midday (between 12:00 h and 14:00 h) to observe crop responses under higher T_air_ variation. These can be considered moderate heatwaves or heat shocks due to the short duration of extreme midday T_air_ (Jagadish et al., 2021; Schulze et al., 2019). They ranged between 32 and 38°C based on the maximum historical T_air_ between 2000 and 2016 for each growth stage. The rest of the days are referred to as “normal days”, following Beijing climatology with midday T_air_ between 20 and 32°C. Plants in each lysimeter were subjected to two surface soil moisture scenarios: i) Wet/control soil (100% surface field capacity, SFC) and ii) Dry soil (40% SFC). Drip irrigation was manually regulated through irrigation (Figure 1). To control soil moisture status and schedule irrigation, soil water content was measured every two to four days with a portable TDR probe (Field Scout TDR 300 portable moisture meter, Spectrum Technologies Inc., Plainfield, IL, USA) at nine locations in each lysimeter. Table S1 shows days after sowing (DAS) when irrigation and chemical fertilizers were applied.

### 2.2. Data acquisition

A small weather station was installed inside the chamber between the two lysimeters (Figure 1). Sensors measuring environmental parameters were connected to a CR-1000 datalogger (Campbell Scientific Inc., Logan, UT, USA), and soil sensors, thermocouples, and photosynthetic active radiation (PAR) devices were connected to a D500 datataker (Pty Ltd, Rowville, Australia). Information was recorded every 10 minutes during the entire crop growing season.

T_air_ and RH were measured using HMP155 HUMICAP Humidity and Temperature Probe (Vaisala Oyj, Vantaa, Finland), wind speed and direction using 034B Wind Sensor (Met One Instruments Inc., Grants Pass, OR, USA) and incoming PAR reached by the canopy and transmitted PAR reaching the soil using SQ-500 Apogee quantum sensor (Apogee Instruments Inc., Logan, UT, USA) (Figure 1). Figure 2A shows diurnal mean and maximum T_air_, mean VPD, mean actual vapor pressure (e_air_) and mean saturated vapor pressure (e*) over the vegetative and reproductive growth stages.

**Figure 2.**
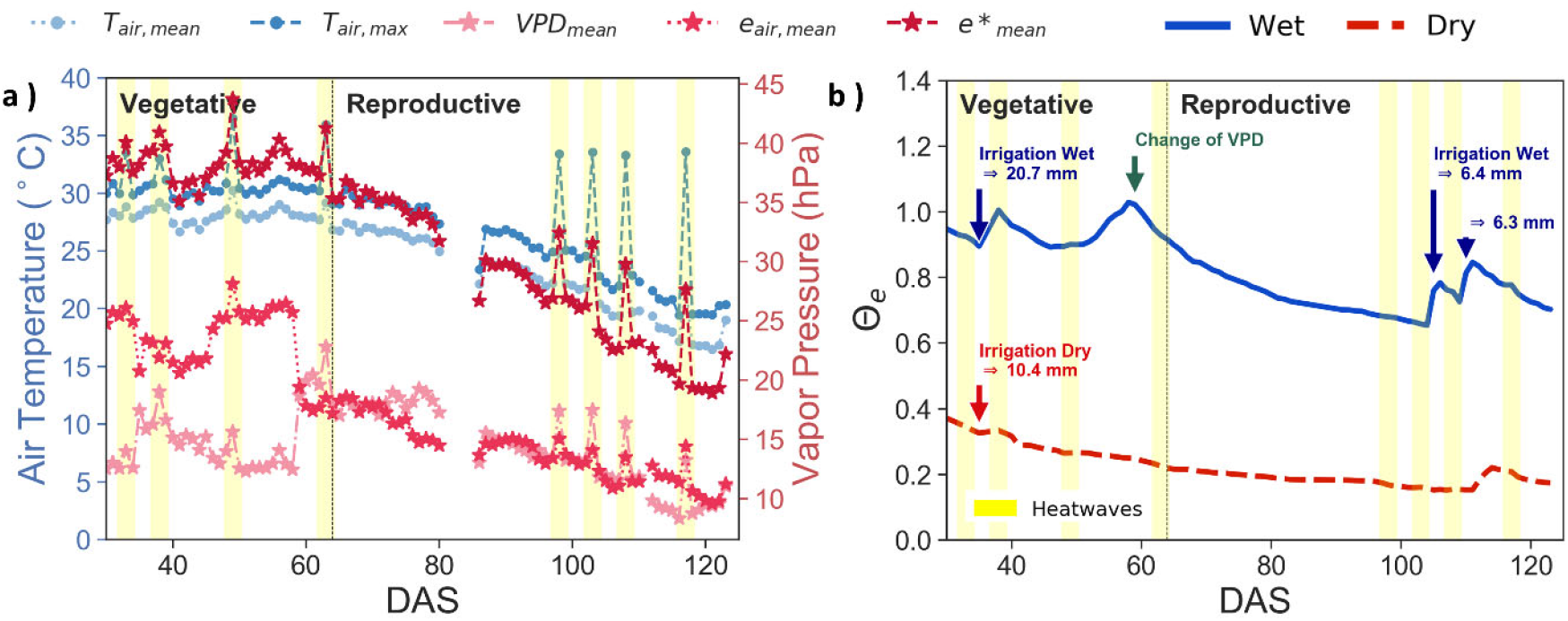
Environmental conditions over the growing season. Yellow vertical lines indicate hot days or moderate heatwaves of one day duration. a) The left axis show mean T_air_, (light blue dots and dotted line), maximum T_air_, (dark blue dots and dashed line), the right axis shows vapor pressure deficit (VPD, light pink starts and dot-dashed line), actual vapor pressure (e = dark pink stars and dotted line) and saturated vapor pressure (e* = red stars and dashed line). b) Daily mean effective soil water content (ϴ_e_) of wet (blue dots and continuous line) and dry (red starts and dashed line) lysimeters. Irrigation amounts are indicated as well as the day that VPD change. DAS = days after sowing.

In each lysimeter, the volumetric soil water content (swc) and soil temperature (T_s_) were measured at the surface in two locations using CS655 Water Content Reflectometer (Campbell Scientific Inc., Logan, UT, USA) and at 20cm depth in one location using CS616 Water Content Reflectometers (Campbell Scientific Inc., Logan, UT, USA). The soil water content reflectometers were calibrated before the experiment with a portable TDR probe by a gravimetric method. To characterize relative soil moisture, we calculated the effective saturation (*ϴ*_e_) as follows:

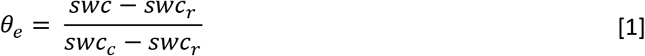

where *swc_r_* is the residual soil water content equal to 2.66% (Chen et al., 2019) and *swc_c_* is the soil water content at SFC (=30%). Figure 2B shows ϴ_e_ at surface layers (up to 20cm depth). In the wet soil lysimeter, ϴ_e_ was between 0.7 and 1, while in the dry soil lysimeter, ϴ_e_ was kept between 0.2 and 0.4.

Evapotranspiration (ET, mm day^−1^, including soil evaporation and canopy transpiration) from each lysimeter was obtained from a water balance approach using irrigation and the water storage change. The later was calculated from the difference between daily average weight measurements over time in each lysimeter. ET corrections were applied to remove noise caused by stabilization after irrigation and other systematic errors (Chen et al., 2019; Tan et al., 2017). Daily average ET from the lysimeters increased after irrigation (Figure S1). ET from the wet soil scenario presented a seasonal average of 1.66 mm day^−1^, significantly higher than ET in the dry soil scenario, with a seasonal average of 1.45 mm day^−1^ and consistent with the total amount of irrigation applied to each lysimeter: 100 mm in wet and 29.5 mm in dry (Table S1). Crop yield was estimated after harvesting to assess productivity differences between the two soil moisture scenarios.

In each lysimeter, four RS PRO Type T thermocouples (2mm Diameter, RS Components Ltd., Corby, Northants, UK) were attached under four leaves with a transpirable tape (see Buchner et al., 2013) for details) to measure T_L_. Thermocouples were calibrated before and after the experiment with a thermal bath following the calibration guide for thermocouples EURAMET cg-8 Version 2.1 (10/2011) (EURAMET, 2011). The water temperature was varied from 0 to 60 °C and measured by a mercury thermometer with 0.1°C accuracy. The temperature sensitivity of each thermocouple was determined and corrected with a linear regression function (R^2^>0.99, RMSE<0.67°C).

Fluorescence techniques measure changes in fluorescence re-emitted from Chl in the photosystems and have been used to assess thermal responses of photosynthesis. Chlorophyll fluorescence parameters were measured at room temperature in three fully extended upper leaves of five plants in each lysimeter, of which four were the same as plants with thermocouples and one was randomly selected. Dark and light-adapted fluorescence measurements with a pulse-amplitude modulation (PAM) Fluorometer (OS5P+ PAM Fluorometer, Opti-Sciences Inc., Tyngsboro, MA, USA) were conducted over 22 days between 11:00 h. and 15:00 h., where seven days were hot days or moderate heatwaves. Before measuring parameters under dark adaptation, leaves were shaded for about 30 minutes. After that, through a saturated light exposure (white light LED with a maximum intensity of 15000 μmols m^−2^ s^−1^), we obtained the maximum fluorescence (F_m_), minimum fluorescence (F_o_), variable fluorescence (F_v_ = F_m_–F_o_) and the F_v_/F_m_ ratio, which estimates the maximum photosynthetic quantum yield of photosystem II photochemistry (Baker, 2008; Humplík et al., 2015).

In addition, after plants were one minute adapted to normal lamp illuminations with a PAR level of around 500 μmol m^−2^s^−1^, the light-adapted fluorescence emission (F’) and the maximal fluorescence (F_m_’) were measured by the PAM. The quantum yield of photochemical energy conversion in photosystem II (Φ_P(II)_) was calculated as:

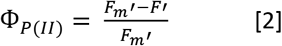

In this study, the sum of the quantum yields of photochemistry (Φ_P(II)_), fluorescence (Φ_F_), constitutive dark-adapted thermal dissipation (Φ_D_) and energy-dependent heat dissipation controlled by mechanisms that regulate the electron transport of photosystems (Φ_N_), is made equal to the unit, as they are mutually exclusive:

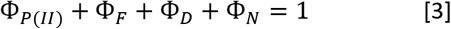

This method was proposed in the Soil Canopy Observation of Photosynthesis and Energy (SCOPE) model (Van der Tol et al., 2014) and applied in other studies (Chen et al., 2019; Lee et al., 2015). We considered the quantum yield of non-photochemical quenching (Φ_NPQ_) as the summed all thermal dissipation (Φ_NPQ_ = Φ_D_ + Φ_N_). Even though this differentiation into two terms is a simplification, it is sufficient for our goals and for preventing equifinality of the model (van der Tol et al., 2014).

Quantum yields can be found in terms of rate coefficients of photochemical reactions (k_p_), chlorophyll fluorescence (k_f_) and heat dissipation from dark-adapted and energy-dependent processes respectively (k_d_+k_n_):

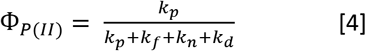

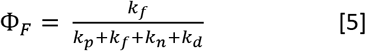

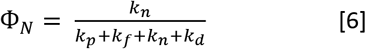

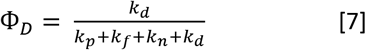

Where k_f_ = 0.05

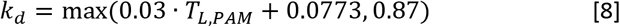

Where T_L,PAM_ is the temperature measured by the PAM at the time of measurements. The optimum temperature T_opt_ was determined as the leaf temperature at which Φ_*P*(*II*)_ reached maximum (see Statistical Analyses below).

The rate coefficients k_N_ and k_p_ vary with the metabolic state, and can be defined by reference to their respective maximum values and considering that k_p_ is (by definition) zero at F_m_’ and at *F_m_* and that k_N_ is zero at *F_m_*.

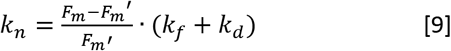

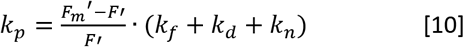

### 2.3. Leaf energy balance model

Leaf stomatal conductance (g_s_) and transpiration (T_r_) were calculated from leaf energy balance fluxes, assuming steady state conditions and neglecting heat storage change and metabolic heat production relative to the other terms (Campbell and Norman, 1998) resulting in Equation 11:

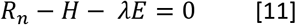

Where R_n_ is the net radiation (W m^−2^), H is sensible heat flux (W m^−2^) and λE is latent heat flux (W m^−2^).

The net radiation flux was obtained as in Campbell and Norman (1998) as follows:

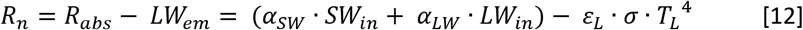

Where R_abs_ is the leaf absorbed radiation (W m^−2^), LW_em_ is the longwave radiation emitted by the leaf (Wm^−2^), α_SW_ and α_LW_ are shortwave and longwave leaf absorptivity (dimensionless), respectively, SW_in_ is the incoming shortwave radiation (Wm^−2^), LW_in_ is incoming longwave radiation (Wm^−2^), ε_L_ leaf emissivity, σ is the Stefan-Boltzman constant (5.6710^−8^ Wm^−2^K^−4^).

α_SW_ was estimated from the albedo based on outgoing and incoming radiance measured with a handheld spectroradiometer (ASD, FieldSpec HandHeld 2™, Analytical Spectral Devices, Inc., Boulder, USA) for 52 samples (above three leaves in four plants in each lysimeter over 13 days) over time, and assumed constant over the whole growing period. The albedo was 0.23 and 0.21 for plants in wet and dry soil respectively with results in the range for maize according to Campbell and Norman (1998). SW_in_ was estimated continuously from incoming PAR measurements using the incoming radiance measurements from ASD to find a scaling relationship with incoming PAR for our relatively constant indoor illumination conditions.

Leaf emissivity (ε_L_) was estimated based on Stefan-Boltzmann law using leaf averaged radiometric brightness temperature measurements from 144 samples (18 dates x 4 leaves x 2 soil scenarios) taken during the growing season with an infrared radiometrically calibrated camera (Fluke Ti55 Infrared FlexCam Thermal Imager, Fluke Corporation, Everett, WA) and T_L_ was measured with thermocouples. For simplicity, the average ε_L_ = 0.98 was used for the whole growing period. Based on Kirchhoff’s law of thermal radiation, *a_LW_ = ε_L_*.

We considered the ground and the walls as LW_in_ (W m^−2^) sources (Monje and Bugbee, 2019), assuming that walls’ temperature was the same as T_air_ and that 80% of the longwave fraction came from the walls, while 20% from the ground with temperature T_s_ (Page et al., 2018):

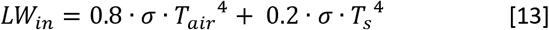

Sensible heat flux, H, (Wm^−2^) was estimated based on the leaf-air thermal gradient (K) based on the mass-transfer equation (Campbell and Norman, 1998):

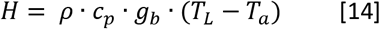

Where *ρ* is the air density (1.292 kg m^−3^), c_p_ is the specific heat capacity of air (1010 J kg^−1^ K^−1^) and g_b_ is the boundary layer conductance to heat (mol m^−2^s^−1^). Assuming negligible the wind inside the growth chamber, g_b_ (mol m^−2^s^−1^) in free convection was based on Campbell and Norman (1998):

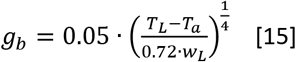

Where w_L_ is the maximum leaf width (m), estimated using average measurements of all leaves of 6 random plants at each lysimeter once a week (13 times), interpolated for continuous values with polynomial regression (R^2^>0.84, data not shown). g_b_ was converted into ms^−1^.

The latent heat flux (λE) is driven by the vapor pressure deficit as in the mass transfer equation:

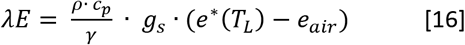

Where g_s_ is total stomatal conductance (m s^−1^), ϒ is the psychrometric constant (hPa K^−1^), e*(T_L_) is the saturated vapor pressure at the leaf (hPa), and e_air_ is the actual vapor pressure of air (hPa).

Inverting the leaf energy balance, g_s_ (mol m^−2^ s^−1^) was obtained as:

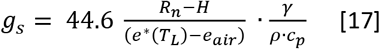

T_r_ (mmol m^−2^ s^−1^) was estimated from the λE, as the mass equivalent using the latent heat of vaporization (λ) and the equivalent mole fraction of water vapor. The evaporative fraction (EF) was estimated as the λE and R_n_ ratio.

In this study, we estimated a new thermal parameter to account for leaf thermal emission excluding the NPQ contribution, referred as “leaf temperature without non-photochemical quenching” (T_L,NPQ=0_). The new slope *β*’ between T_L,NPQ=0_ vs T_air_ would be an indicator of the thermoregulation excluding the effect of thermal regulatory dissipation. To estimate T_L,NPQ=0_, the energy of thermal emission associated with NPQ (Φ*_NPQ_* · *PAR_abs_*) was subtracted from the LW_out_ using Stefan-Boltzman law [Eq. 4]:

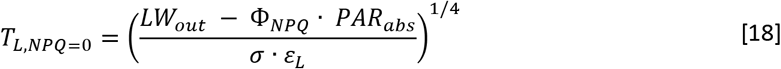

Where PAR_abs_ is the absorbed PAR obtained from incoming PAR multiplied by the shortwave absorptivity and σ is the Stefan-Boltzman constant (= 5.67E-8 W m^−2^ K^−4^).

### 2.4. Statistical Analysis

Instantaneous T_L_ measurements between 6:00 h. and 19:00 h. and between 12:00 to 14:00 h. were used to evaluate *β* and plant responses to T_air_. The coefficient of variation (cv) of T_air_ and T_L_ was calculated for diel, diurnal, midday, and nocturnal measurements as an indicator of range of variability in temperatures. The thermal stress or thermal shock (T_shock_) was estimated as the difference between maximum and minimum diurnal T_air_.

For fluorescence measurements, the mean value and standard error of replicas in each lysimeter were taken. We applied ordinary least squares linear and polynomial regression to assess relationships among variables throughout study. We only used second order polynomial relationship for fluorescence variables versus temperatures. T_opt_ was evaluated as the maximum of the second order polynomial relationship between T_L_ and Φ_P(II)_ as maize is a C4 plant (Crafts-Brander and Salvucci et al., 2002).

To assess mean differences between hot days and normal days and dry and wet treatments the Wilcoxon test (non-parametric) with Bonferroni correction was applied as it allows comparisons without any assumption about data distribution for dependent variables. We used the coefficient of determination (R^2^), upper and lower 95% confidence interval (CI) values and p-value with a significant level of 5% (*p ≤ 0.05, **p ≤ 0.01, *** p ≤ 0.001 and ns = no significant) to evaluate regressions (linear and polynomial) among variables.

A two-way analysis of variance (ANOVA) was used to investigate whether significant changes for several parameters were occurring between the two soil moisture scenarios and between normal days with heatwaves. We compared means from all the instantaneous measurements between 6:00 h. and 19:00 h. over the season and also from parameters just at the time of fluorescence measurements using the Python package Pingouin (Vallat, 2018).

## 3. Results

### 3.1. Maize thermoregulation under wet and dry soil scenarios

Maize plants operated within a narrower T_L_ range than the air T_air_ during diurnal hours, especially for the wet scenario and midday as shown by the coefficient of variation (cv) in Table 1. In contrast, the variability of diel T_L_ (24h) was slightly higher than T_air_ variability for the wet scenario and quite similar for the dry scenario. At night the situation was similar with a larger variation in T_L_ than T_air_. Thus, the lower diel cv in T_L_ than T_air_ is entirely due to the influence of night conditions.

**Table 1.** Coefficient of variation (cv) of air temperature (T_air_) and leaf temperature (T_L_) of wet and dry soil moisture scenarios measured every 10 minutes with total (24h) (n = 2496 observations), diurnal (between 6:00 h. and 19:00 h), nocturnal (between 19:00 h. and 6:00 h.) and midday (between 12:00 h. and 14:00 h.) averages during the growing season.

| **Period** | **cv $T_{air}$** | **cv $T_{L,Wet}$** | **cv $T_{L,Dry}$** |
| --- | --- | --- | --- |
| **Total (Diel)** | 0.213 | 0.229 | 0.218 |
| **Diurnal** | 0.172 | 0.128 | 0.133 |
| **Midday** | 0.148 | 0.115 | 0.119 |
| **Nocturnal** | 0.197 | 0.216 | 0.207 |

Observations of T_air_ versus T_L_ (Figure 3) showed that T_L_ was significantly correlated with T_air_ (r^2^>0.77, p<0.001) in both soil scenarios. T_L_ got closer to T_air_ at warmer temperatures, reducing ΔT, which indicates partial thermoregulation (Michaletz et al., 2016). This is shown by the least-squares regression fitted slope of 0.71 for wet soil and 0.76 for dry soil scenario. T_eq_ was found from the regression lines intercepting the 1:1 line at around 35°C and 38°C for wet and dry soil respectively. Depending on the aggregation and time of the day of different data subsets, T_eq_ could vary between 33°C and 38°C for the wet soil (Figure S2) but in all instances, T_eq_ was about 2 to 3°C higher in dry soil scenario than in the wet, with T_air_-T_L_ slopes between 0.65 and 0.8.

**Figure 3.**
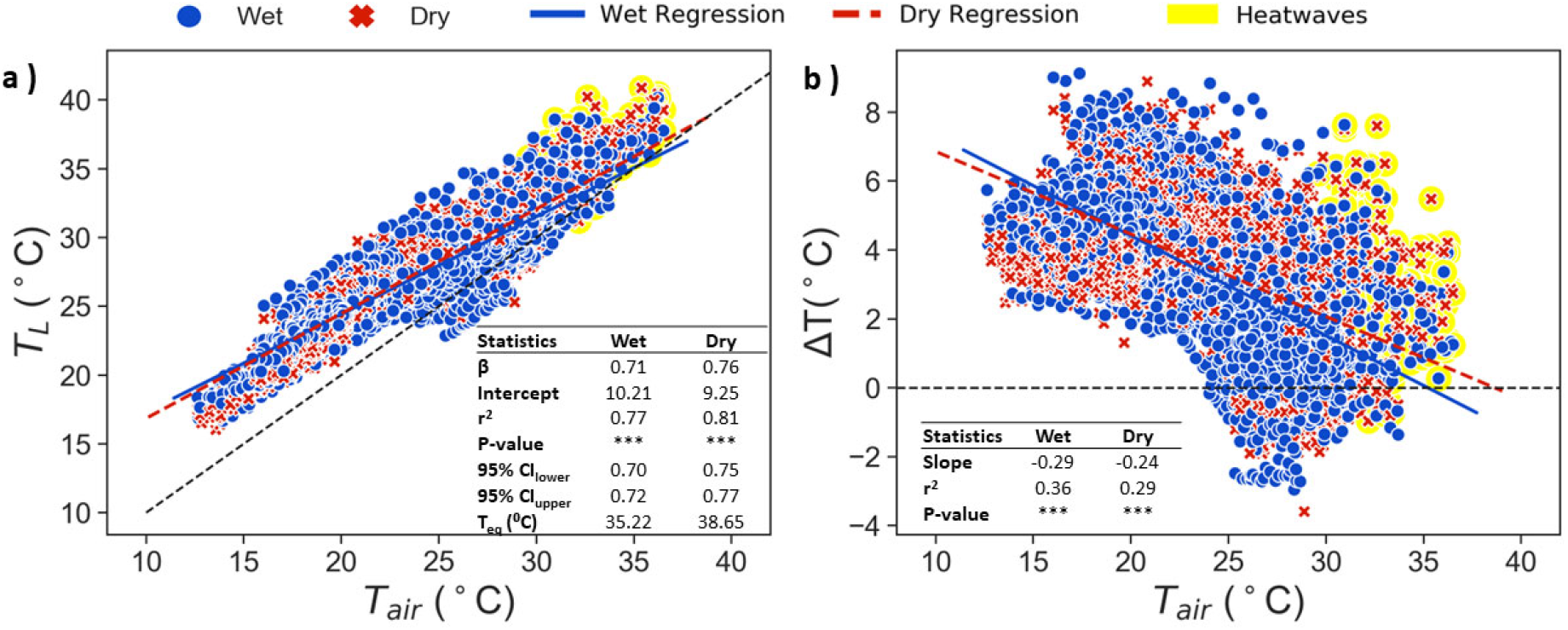
a) Air temperature (T_air_) vs leaf temperature measured by thermocouples (T_L_). b) T_air_ vs leaf temperature excess (ΔT) for instantaneous measurements between 6:00 a.m. and 7:00 p.m. for wet (blue dots) and dry soil (red crosses). First order regression lines are shown. Table shows the slope (β), intercept, coefficient of determination (R^2^), p value (Wilcoxon test with Bonferroni correction, *p <= 0.05, **p <= 0.01, *** p <= 0.001 and ns = no significant), upper and lower 95% confidence interval (CI) values and the leaf-air equivalence temperature (T_eq_), where T_air_=T_L_. based on regression line. Maximum T_air_ during heatwaves are shaded in yellow.

### 3.2. Thermoregulation mechanisms during the growing season: NPQ and stomatal conductance

We evaluated if thermal dissipation from NPQ contributed to thermoregulation using the photochemistry dataset. To support our analysis, we assessed first whether thermoregulation responses established from air and leaf temperature observations taken at the time of the photochemistry dataset were comparable to the ones from the continuous dataset (Figure 3). In fact, results from the relationship between T_air_ vs T_L_ were quite similar (Figure 4A) to those from Figure 3 indicating partial thermoregulation or limited homeothermy, with T_eq_ at around 38°C and 41°C for wet and dry soils, respectively and negative ΔT slopes in relation to T_air_ (Figure 4C).

**Figure 4.**
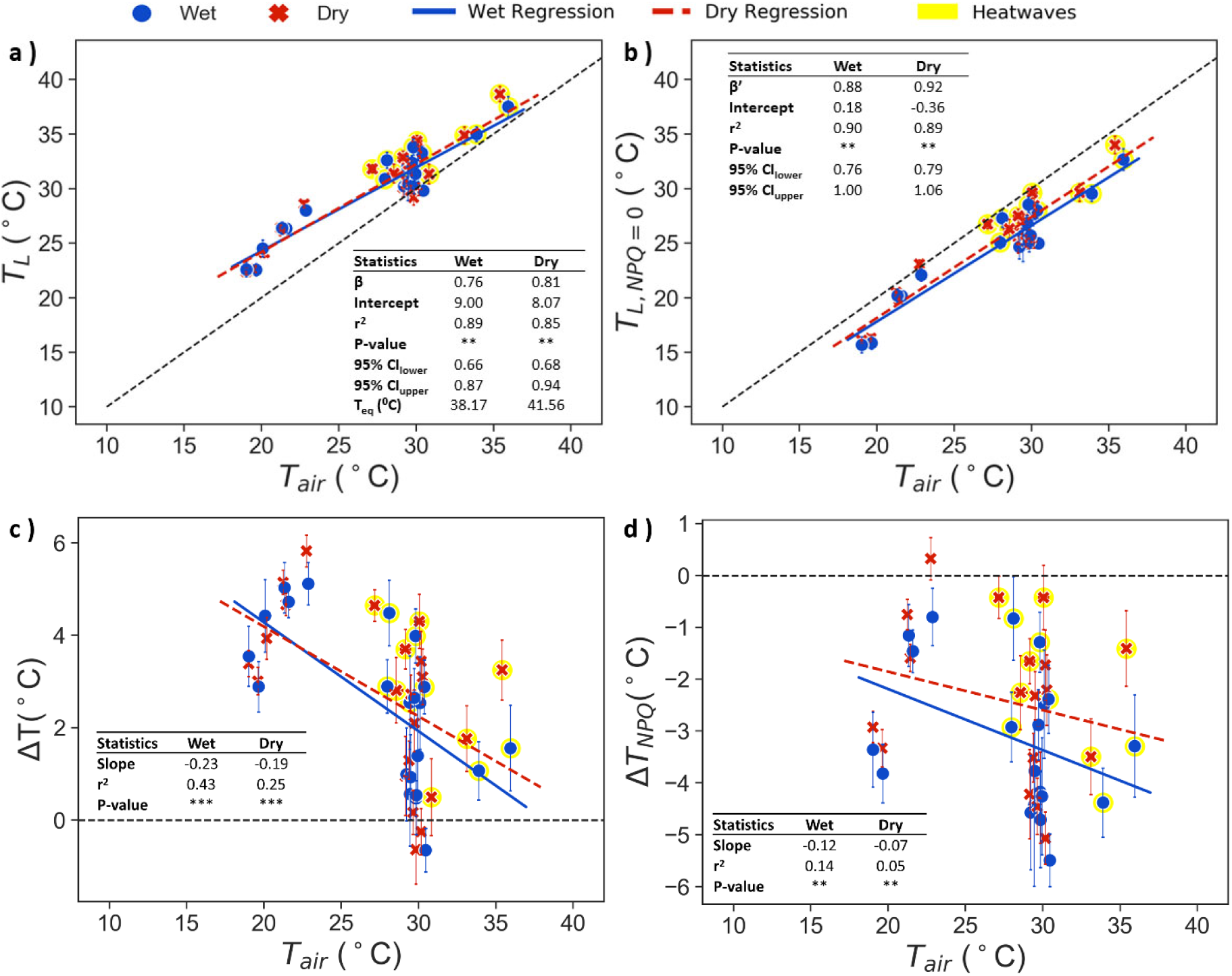
Air temperature (T_air_) vs a) leaf temperature (T_L_), b) leaf temperature corrected for NPQ emission (T_L,NPQ=0_) at the time of fluorescence measurements (around midday), c) leaf temperature excess (ΔT) and d) leaf temperature excess calculated with the T_L,NPQ=0_ (ΔT_NPQ_) of wet soil in blue dots and dry soil scenario in red crosses. Continuous blue line and dashed red line show first order regression for wet and dry data, respectively. Table inception shows the slope (β or β’), intercept, coefficient of determination (R^2^), p value (Wilcoxon test with Bonferroni correction, *p <= 0.05, **p <= 0.01, *** p <= 0.001 and ns = no significant), upper and lower 95% confidence interval (CI) values and the leaf-air equivalence temperature (T_eq_), where T_air_=T_L_ based on regression line. Moderate heatwaves during midday are shaded in yellow.

It is remarkable that this partial thermoregulation response disappeared when we accounted for the contribution of NPQ to heat dissipation, getting closer to poikilothermy. We subtracted from T_L_ the longwave energy equivalent from the thermal dissipation from NPQ, obtaining T_L,NPQ=0_ (Figure 4B). T_L,NPQ=0_ vs. T_air_ showed a slope closer to 1 especially for the dry soil scenario (slope 0.92) and a slope closer to 0 compared to ΔT_NPQ_ (Figure 4D). Remaining variations in T_L_ after controlling for NPQ, e.g. in T_L,NPQ=0_ can be attributed to variations in other leaf energy balance terms like the cooling from T_r_ but in any case show a tighter coupling to T_air_ (slopes being closer to 1) than with original T_L_.

From these analyses, NPQ dissipation can be identified as a negative feedback mechanism from photosynthesis supporting leaf thermoregulation towards T_opt_, helping plants to maintain narrower T_L_ ranges in response to T_air_ variations (Figure 4). Thus, the further away T_air_ was from T_opt_, the higher the thermal dissipation from NPQ, which results in T_L_ increasing above T_air_. The heat dissipation from NPQ was minimum around T_opt_, where Φ_P(II)_ was maximum, with T_L_ being in equilibrium with T_air_ (T_eq_). Even though the energy contribution from NPQ emission to average LW_out_ (Wm^−2^) was only around 7% (equivalent to 5.85°C) in our experimental conditions, it was enough to help plants to thermoregulate in suboptimal conditions, below T_opt_. Dissipation of thermal energy from NPQ represents about 53% of the incoming PAR and 82% of absorbed PAR. Our experiment does not allow to establish what happens when T_L_ is above T_opt_ as the temperature range of the photochemical dataset, following the climatology in Beijing without causing lethal damage, was only slightly above the T_opt_ for maize.

To assess if the T_eq_ determined from Figure 4 was close to the T_opt_, T_opt_ was estimated as the T_L_ for maximum Φ_P(II)_.

Figure **5** shows that Φ_P(II)_ increased non-linearly with T_L_ following a second-order polynomial regression with R^2^ = 0.66 (p<0.001) for the wet and R^2^ = 0.43 (p<0.001) for the dry scenario. While Φ_P(II)_ increased with T_L_, Φ_NPQ_ decreased due to the trade-off between Φ_P(II)_ and Φ_NPQ_ (linear correlation R^2^>0.98, p<0.001 in Figure S3). From the Φ_P(II)_ vs. T_L_ curve, the T_opt_ identified at around 38°C and 41°C for wet and dry scenarios respectively (

**Figure 5.**
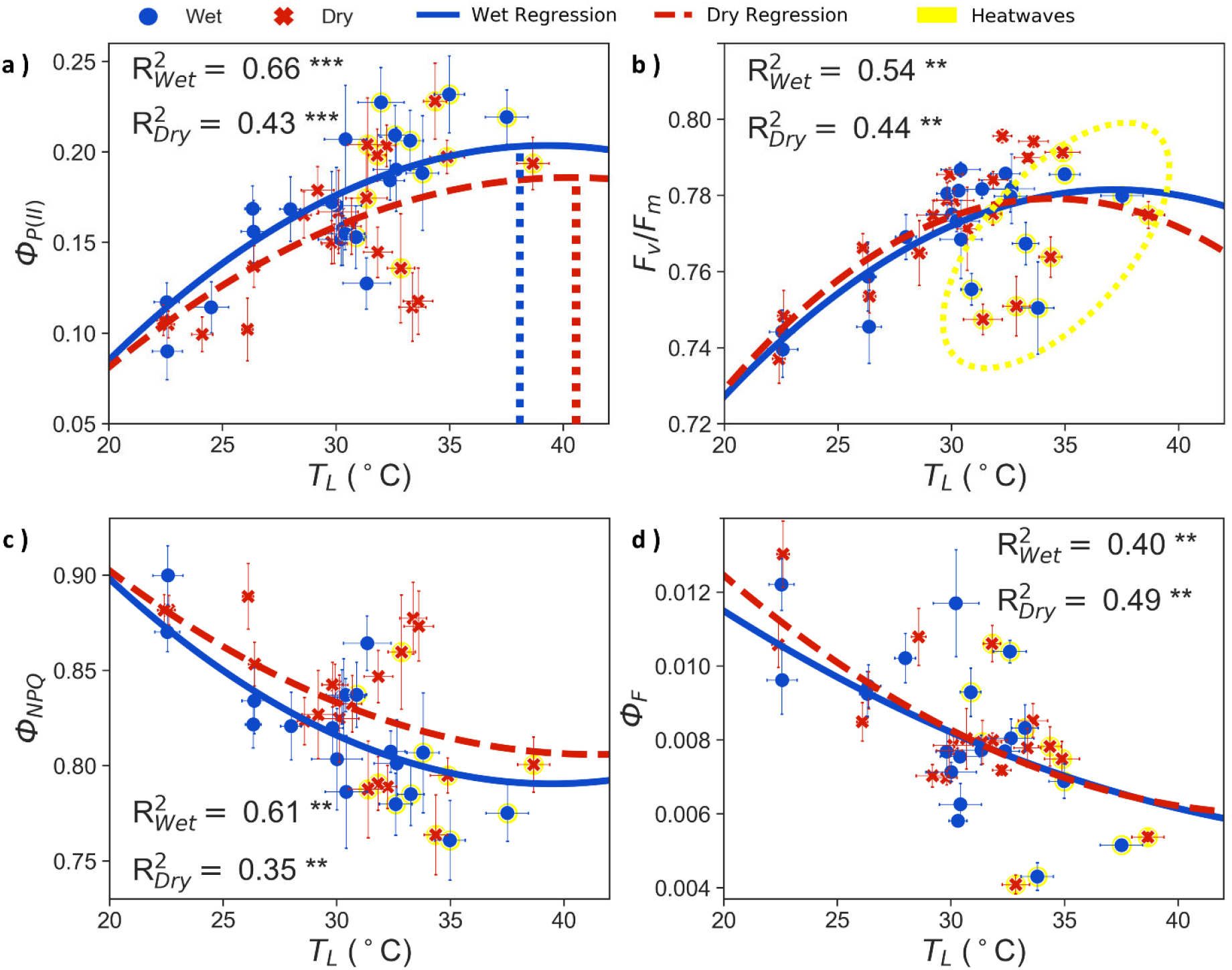
a) Quantum yield of photochemistry (Φ_P(II)_), b) Maximum quantum yield of photosystem II photochemistry (F_v_/F_m_), c) quantum yield of non-photochemical quenching (Φ_NPQ_) and d) quantum yield of fluorescence (Φ_F_) compared to T_L_ measured at the time of the fluorescence observations for wet (blue dots) and dry (red crosses) scenarios. Error bars show the variation among replicas. R^2^ and p value (Wilcoxon test with Bonferroni correction, *p <= 0.05, **p <= 0.01, *** p <= 0.001 and ns = no significant) are given for wet and dry soil data from a second order polynomial regression for wet (continues blue line) and dry (dashed red line). Moderate heatwaves are yellow shaded and circled. Maximum Φ_P(II)_ is identified with vertical lines associated to optimal temperature for photosynthesis (T_opt_).

Figure **5**A), matched well the T_eq_ from Figure 4A (38.2°C and 41.2°C). This highlights the role of thermoregulation as an adaptive mechanism to optimize photosynthesis around T_opt_, as Φ_P(II)_ in C4 plants is a good surrogate for photosynthesis Crafts-Brander and Salvucci et al., (2002).

The increase in Φ_P(II)_ (and correlative decrease in Φ_NPQ_) with T_L_ was slightly higher for plants in the wet than in the dry soil (around 13% and 10% respectively) indicating better photosynthetic efficiency with lower heat dissipation for wet soil conditions (

Figure **5**C). Φ_F_ also decreased towards T_opt_ in both soil scenarios (R^2^ > 0.40, p<0.01) but in the case of dry soil, Φ_F_ showed the best correlations with increasing T_L_ than with other photochemical variables (

Figure **5**D). Although significant, the relationship of Φ_F_ and Φ_P(II)_ is much weaker than between Φ_NPQ_ and Φ_P(II)_ (R^2^=0.98-1), with a negative, moderate trend only for wet soil (Figure S3). Another key indicator of photochemical responses to temperature is F_v_/F_m_. In our conditions, F_v_/F_m_ increased with T_L_, peaking at a lower T_L_ (around 34 °C) than Φ_P(II)_, especially for the dry soil.

Stomata contribute as well to thermoregulation as plants have to balance carbon gains and water losses responding to changes in the efficiency of photosynthesis. This is shown in Figure 6 where g_s_ and T_r_, directly derived from measured T_L_ (limited homeothermy) were always lower than for the hypothetical case of TL=T_air_ (poikilothermy), which assumed that stomata are regulated to maximize cooling and T_r_, maintaining the evaporative fraction (EF) around 1. Stomatal regulation dynamics under limited homeothermy to VPD are also more coherent than just to T_L_, showing an optimum of around 14hPa in both soil scenarios and a second minimum at higher VPDs (> 35 hPa). However, when assuming a poikilothermic thermoregulation, g_s_ decreased exponentially with increasing VPD (Figure 6E and F).

**Figure 6.**
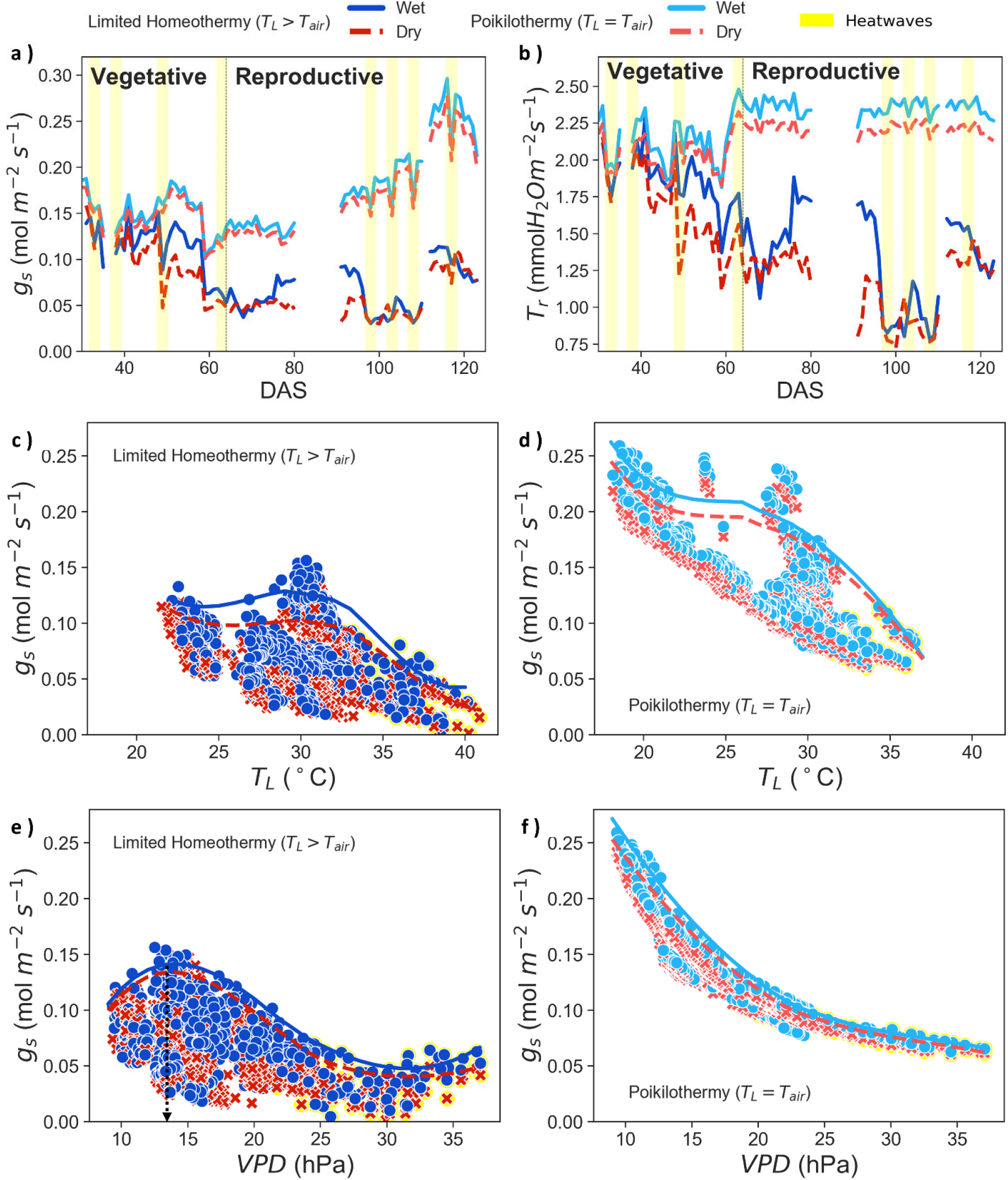
a) Diurnal mean stomatal conductance (g_s_) and b) diurnal mean transpiration rates (T_r_) over the growing season for measured T_L_ (limited homeothermy) of wet (dots and blue continuous line) and dry (crosses and red dashed line) scenarios and for simulated poikilothermy (T_L_=T_air_) of wet (light blue continuous line and dry (light red dashed line). DAS = Days after showing. c) g_s_ vs T_L_ for limited homeothermy, d) g_s_ vs T_L_ for poikilothermy, e) g_s_ vs vapor pressure deficit (VPD) for limited homeothermy and e) g_s_ vs VPD for poikilothermy with VPD threshold around 14 hPa (arrow indication). Moderate heatwaves are yellow shaded.

The role of soil water availability in thermoregulation was clear as for any T_L_ range, the overall g_s_ of the dry soil scenario was lower than that of wet soil, reflected also in an overall lower T_r_ and cooling capacity (lower EF) (Figure 6 and Table 2). Nonetheless, dry soil g_s_ presented the same relative patterns as wet soil in response to VPD (Figure 6). This lower g_s_ reduces CO_2_ uptake impacting crop lifetime carbon gain and final yield which was significantly lower in the dry soil condition (Table 2). The higher T_opt_ and T_eq_ in dry soil plants suggest acclimation to the drier condition but at the cost of achieving lower photochemical efficiency, with reduced T_r_ and suboptimal g_s_ that impair CO_2_ uptake ultimately affecting crop yield (Table 2).

**Table 2.** Mean differences between plants grown in wet or dry soil (Water Status) and for observations corresponding to average air temperature days compared to hot days (Heat condition) for instantaneous observations and for the subset corresponding to the time of fluorescence observations.

| Parameters | Instantaneous (n = 2496) |  | Fluorescence (n=24) |  |
| --- | --- | --- | --- | --- |
|  | Water Status | Heat Condition | Water Status | Heat Condition |
| $T_{air}$ (°C) | ns | ***↑ | ns | ***↑ |
| $T_L$ (°C) | ns | ***↑ | ns | ***↑ |
| $\Delta T$ (°C) | ns | ns | ns | ns |
| $T_{shock}$ (°C) | ns | ***↑ | ns | ***↑ |
| $g_s$ (mol m<sup>-2</sup> s<sup>-1</sup>) | ***↓ | ***↓ | ns | *↓ |
| $T_r$ (mmolH<sub>2</sub>O m<sup>-2</sup> s<sup>-1</sup>) | ***↓ | ns | ns | ns |
| EF | ***↓ | ns | ns | ns |
| ET (mm day<sup>-1</sup>) | **↓ | ns | ns | ns |
| $\Theta$ | ***↓ | ns | ***↓ | ns |
| $\Phi_{P(II)}$ | | | ns | **↑ |
| $\Phi_F$ | | | ns | ns |
| $\Phi_{NPQ}$ | | | ns | **↓ |
| $F_v/F_m$ | | | ns | ns |
| Yield (kg ha<sup>-1</sup>) |  |  | *↓ |  |

Air temperature ( $T_{air}$ ), leaf temperature ( $T_L$ ), leaf-air temperature difference ( $\Delta T$ ), diurnal $T_{air}$ oscillation ( $T_{shock}$ ), stomatal conductance ( $g_s$ ), transpiration rate ( $T_r$ ), evaporative fraction (EF), evapotranspiration (ET) and relative soil water content ( $\Theta$ ) over the season, quantum yields of photosynthesis ( $\Phi_{P(II)}$ ), fluorescence ( $\Phi_F$ ), non-photochemical quenching ( $\Phi_{NPQ}$ ) and maximum quantum yield of photochemistry II ( $F_v/F_m$ ). Crop yield (yield) at the end of the experiment. Significance level: \*p <= 0.05, \*\*p <= 0.01, \*\*\* p <= 0.001 and ns = no significant. The arrows indicate an increases ↑ or decreases ↓ when comparing values in wet soil plants compared to dry water soil (Water Status) and values for average $T_{air}$ days compared to hot (extreme) days (Heat condition).

Note that ET measured from lysimeters was higher in the wet scenario and normal days, supporting the findings from our leaf T_r_ model, with lower T_r_ for the dry soil scenario than the wet soil despite the scale mismatch between both variables (Table 2).

### 3.3. Thermoregulation mechanisms during hot days: NPQ and stomatal conductance

Compared to normal days, hot days showed significantly lower Φ_NPQ_ and higher Φ_P(II)_ averaged over the season (Table 2) which made sense as T_L_ was generally closer to T_opt_ on those days. Looking at photochemistry responses to T_L_, the strongest indicator of heat stress was F_v_/F_m_ which decreased in hot days while in normal days it increased almost linearly with T_L_ (

Figure **5**B, hot days circled in yellow). This pattern was not clear for Φ_P(II)_ Φ_NPQ_ or Φ_F_ with more scattered observations in hot days than in normal days around the curve (Figure **5**).

Isolating only hot days, F_v_/F_m_ was clearly sensitive to the sudden temperature rise of hot days, decreasing with T_shock_ (R^2^=0.76) for both soil moisture scenarios (Figure 7). However, only a weak negative correlation with T_shock_ was found for Φ_P(II)_ (R^2^=0.15) and a positive one for Φ_NPQ_ (R^2^=0.20). This trend was just due to plants in the wet soil, which increased (decreased) about 30% their Φ_NPQ_ (Φ_P(II)_) with a 10°C increase in T_shock_ while Φ_NPQ_ and Φ_P(II)_ of plants in the dry soil, already at a lower level of photochemical efficiency at that point, seem less sensitive to changes in T_shock_. Nonetheless, due to the small sample size, this interpretation of wet and dry soil has to be taken with caution.

**Figure 7.**
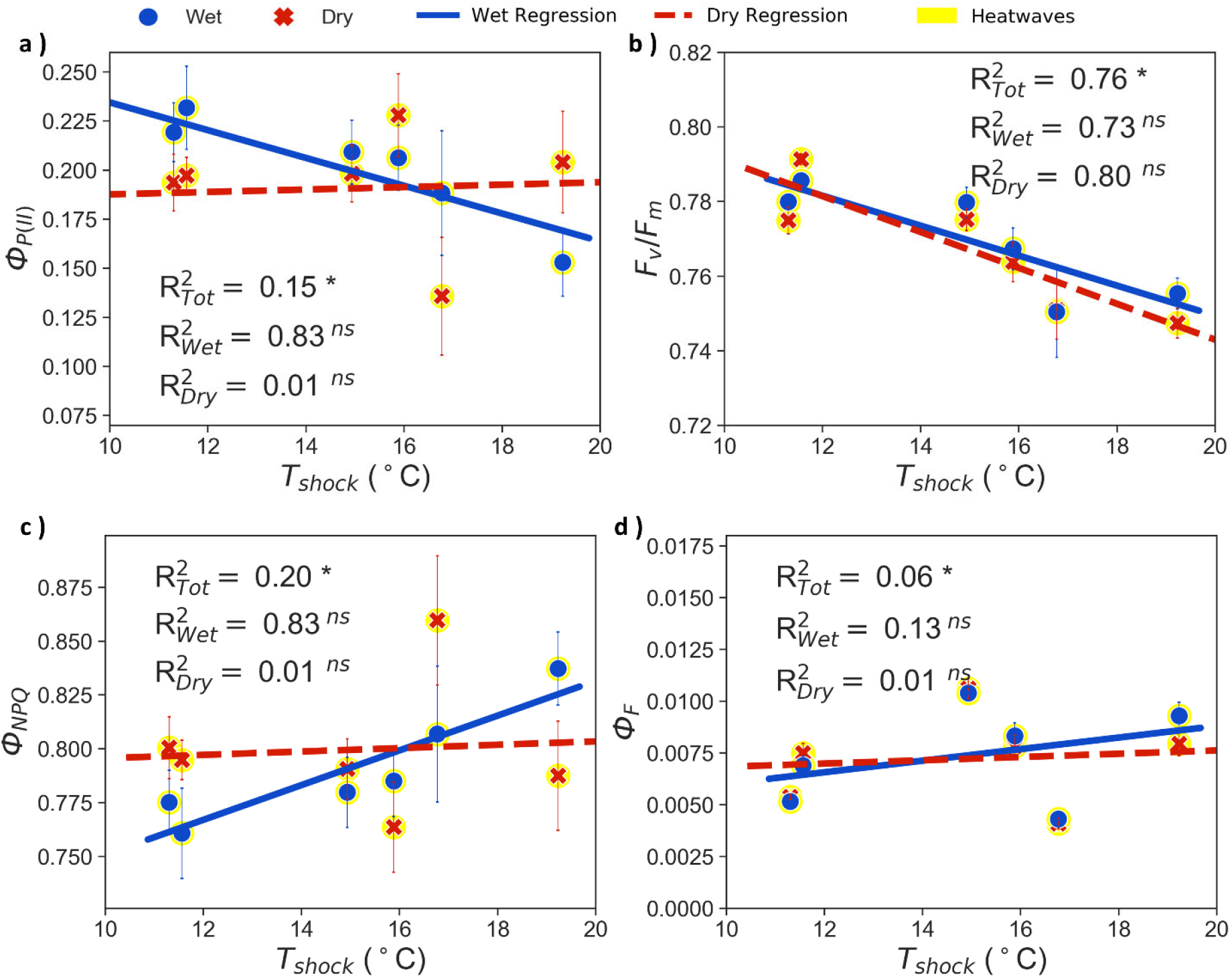
Relationship of T_shock_ (maximum T_air_ diurnal oscillation) during moderate heatwaves with a) quantum yield of photochemistry (Φ_P(II)_), b) maximum quantum yield of photosystem II photochemistry (F_v_/F_m_), c) quantum yield of non-photochemical quenching (Φ_NPQ_) and d) quantum yield of fluorescence (Φ_F_) with linear regression for wet (dots, continuous blue line) and dry (crosses, dashed red line) scenarios. R^2^ and p value (Wilcoxon test with Bonferroni correction, *p <= 0.05, **p <= 0.01, *** p <= 0.001 and ns = no significant) are given for wet scenario, dry scenario and all the data (tot=total). Error bars correspond to 15 samples and 4 replicas in T_L_.

Regarding cooling and stomatal control, hot days showed significantly lower g_s_, and higher T_L_ averaged over the season than normal days, with no significant differences in evaporative cooling (EF), T_r_ or canopy level ET (Table 2). Looking at subdaily patterns it is very clear the larger role of stomatal closure during hot days compared to normal days (Figure 8 and Figure S4). g_s_ followed the diurnal course of temperature, with minimum g_s_ happening at midday corresponding to the temperature peak. Hot days concurrent with dry soil result in an even lower g_s_ reaching almost total closure at midday. Hot days also presented larger hysteretic behavior with asymmetry between morning and afternoon, which was more marked for the dry soil (Figure and Figure S4).

**Figure 8.**
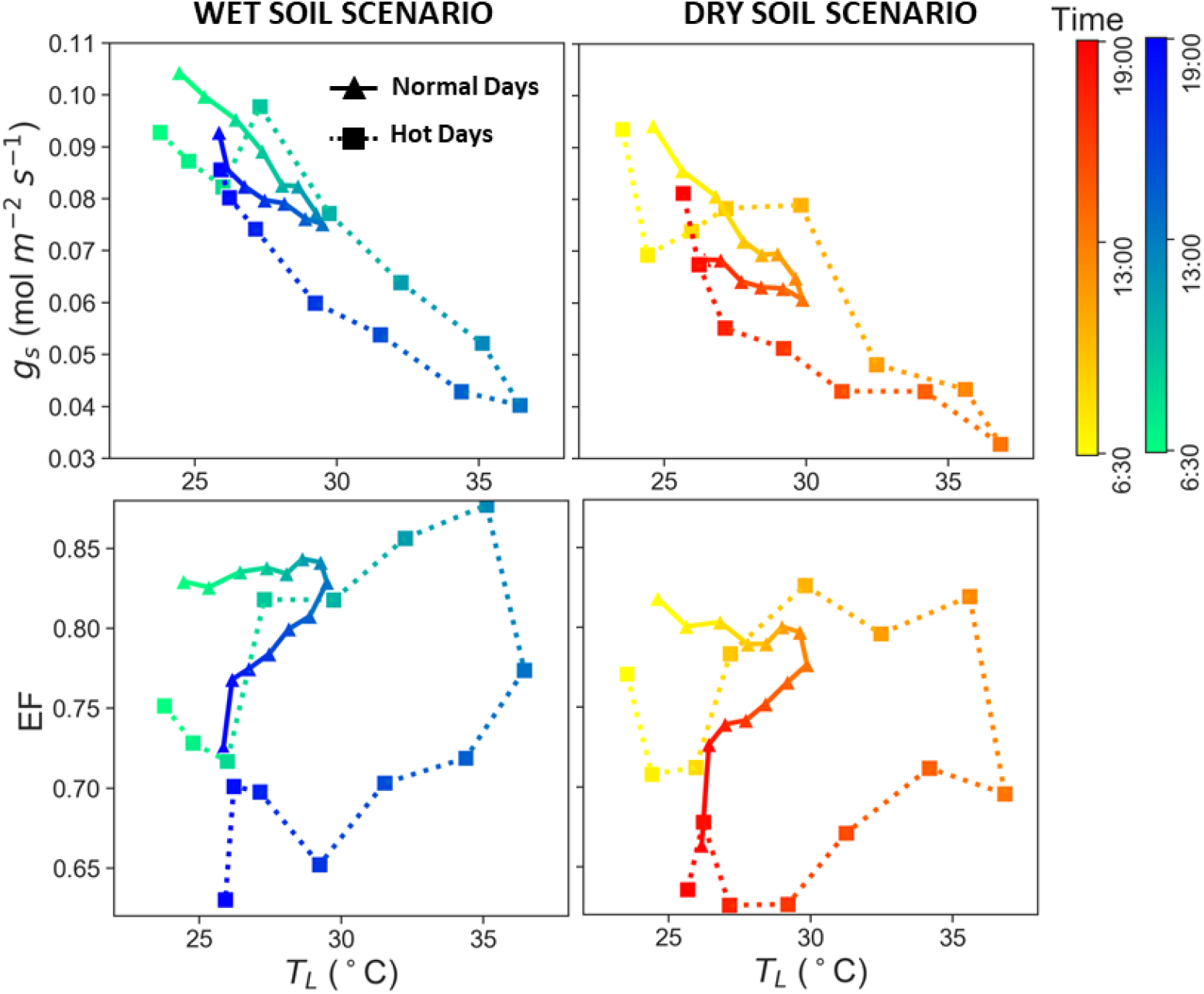
Hysteresis curves showing hourly diurnal evolution between 6:30 a.m. and 7:00 p.m. averaged over the season for stomatal conductance (g_s_) and evaporative fraction (EF) vs. T_L_ for wet (blue-green colormap) and dry (red-yellow colormap) soil moisture scenarios for normal and hot days (heatwaves). Each observation represents the average of hourly observations across the season.

Even though g_s_ was minimum at midday, cooling (EF) was maximum at that time and larger for hot days than normal days but only for the morning hours and only for wet soil (Figure). In the afternoon hours, EF of hot days was always below that of normal days. These differences between wet and dry conditions and between morning and afternoon explain that no significant differences were found with T_r_, EF, and ET averages over the season (Table 2).

Plants in hot days regulated stomata to avoid drying out, but T_r_ is sustained because as VPD is higher, the vapor diffusivity is higher. However, in dry soil conditions their capacity to transpire and cool down is further reduced by a larger stomata closure.

In summary, in hot days, there seem to be two opposite effects taking place. On one hand, as T_L_ becomes closer to T_opt_ the photosynthetic efficiency tends to increase as shown in the ANOVA (Table 2), reducing Φ_NPQ_ and increasing Φ_p(II)_, to a larger extent in the wet treatment where plants can have higher cooling reducing thermal stress and had an overall higher photochemical efficiency. On the other hand, as the magnitude of the thermal shock becomes larger that positive effect of higher temperatures is reversed, as a higher T_shock_ increases Φ_NPQ_ and reduces Φ_p(II)_. Water availability modulates these responses, with plants in hot days and dry soil not being sensitive to changes in T_shock_ as they already had a lower photochemical efficiency.

## 4. Discussion

Understanding maize thermoregulation in response to increases in T_air_ is complex as plants are influenced by several factors such as variety, life history and environmental conditions (Schulze et al., 2019; Yamori et al., 2014), including decreases in soil water availability. In natural conditions, covariation of T_air_ and irradiance makes it difficult to separate photochemical and stomatal responses to thermal changes from those to irradiance changes. By controlling some of the environmental conditions in a growth chamber, with constant illumination, it is possible to isolate leave responses to seasonal and diurnal changes in T_air_ (Roy et al., 2021).

### Limited homeothermy evidence

In the experiment, we found evidence supporting the hypothesis that maize regulates T_L_ to maintain stable metabolic function around the photosynthetic optimum. The T_opt_ estimated by the optimum in the relation between Φ_P(II)_ and T_L_ was close to the T_eq_ (

Figure **5**), supporting that thermoregulation aims elevate T_L_ at around T_opt_, being the T_eq_ a good estimate of T_opt_ (Figure 3 and Figure S2). Our findings for T_eq_ are in the range for T_opt_ for maize defined by Hatfield et al. (2011), between 33°C and 38°C.

Maize plants during the growing season behaved as limited homeotherm, in line with a study conducted with long-term global temperature data for 64 species. However, in Michaletz et al. (2016) only few C4 observations and from a single genus (*Atriplex*) were included.

We assessed thermoregulation based on the fitted slope or *β*, between T_air_ and T_L_ (0.71 for wet and 0.76 for dry scenario, Figure 3) as in Mitchaletz et al. (2016). Additionally, we also considered lower ranges of variability in T_L_ than in T_air_ during diurnal and midday conditions as a strong indicator for thermoregulation (Table 1).

### Role of non-photochemical quenching on thermoregulation

A key aspect in our study was to not only establish the value of *β*, which is useful to predict rates of photosynthesis, respiration or transpiration in earth system modeling but contribute to developing theory for variations in *β* during climatic average conditions (Blonder and Mitcheletz, 2018).

As T_L_ results from balancing and feedback among various incoming and outgoing energy fluxes, it is complex to isolate individual contributions and particularly NPQ, especially as energy balance models do not incorporate heat dissipation from NPQ even though it can represent up to 80% of absorbed PAR (Demmig-Adams et al., 2006). In our study, we were able to explain variations in *β* by controlling T_L_ for the contribution of heat dissipation from photochemistry (T_L,NPQ=0_). The new slope *β*’ (T_L,NPQ=0_ vs T_air_) became higher and closer to 1 (Figure 4), approaching the behavior of a poikilotherm (T_L,NPQ=0_ ~ T_air_). This provides for the first time indication of biological thermoregulation in suboptimal temperatures by NPQ and photochemistry.

A direct contribution of NPQ to T_L_ was already shown experimentally by Kaňa and Vass (2008) in tobacco leaves. In their study, the higher NPQ happened as a photoprotection response to higher irradiance on distant areas of the leaf. By suppressing evaporative cooling artificially, they showed a direct contribution of NPQ heat dissipation to the longwave outgoing radiation independently of g_s_. In our case, the warming effect of NPQ seen on T_L_ was higher at cooler or suboptimal T_air_ (Figure 4), associated with higher g_s_ (Figure 6) which should actually reduce the excess temperature ΔT. Thus, g_s_ dynamics do not explain the observed *β* in our experiment.

The reason for an increased NPQ is photoprotection from excess solar energy absorbed by the light-harvesting system, removing it safely as thermal energy, to avoid formation of reactive oxygen species (ROS) and affecting also fluorescence quenching (Demmig-Adams et al., 2006). It is activated when environmental conditions restrict plant growth, such as when air temperature is below or above T_opt_, causing intrinsic photosynthetic capacity to drop. There are multiple ways in which thermal energy is dissipated, some are reversible, such as those that happen during hot days, and others are sustained over time, such as those taking place between wet and dry scenarios. In general, important changes in the xanthophyll cycle take place. For reviews of processes behind the dissipation of energy via NPQ refer to Porcar-Castell et al., (2014) and Demmig-Adams & Adams, (2006).

Most of the studies quantifying increases in NPQ due to lower photochemical efficiency are related to increases in solar irradiance. Few studies focused on the effect of suboptimal temperatures on photochemistry and had similar findings to ours, but they did not assess the effect on thermoregulation. For example, Tsonev and Hikosaka (2003) and Hirotsu et al., (2004) found that thermal dissipation increased proportionally to the decrease in Φ_P(II)_ for both *Chenopodium album* and various rice cultivars respectively in response to cooling.

Our findings contribute to understanding plant mechanisms of warming up within the *limited homeothermy* hypothesis proposed by Mahan & Upchurch (1988). They stated that plants are capable of actively decreasing its T_L_ above the T_opt_ by transpiration cooling, while their capability to warm up in cold conditions (T_L_ < T_opt_) is limited, resulting in T_L_ fluctuating with ambient microclimate factors, as respiration has a very marginal effect in plant cells and heliotropism is restricted to few species. Our findings show evidence that in suboptimal conditions heat dissipation from NPQ contributes actively to the warming of T_L_ above T_air_. The high negative correlation between Φ_P(II)_ and Φ_NPQ_ (Figure S3) indicates that limited homeothermy responses due to heat dissipation were associated with lower photosynthetic efficiency below the T_opt_ (Figure **5**) as these are competing processes.

### Role of soil water availability on thermoregulation at seasonal scale

Plants in the wet soil presented slightly a larger capacity to thermoregulate, with lower cv and *β* than those in the wet treatment, with a narrower range in T_L_ than plants in the dry scenario (**Error! Reference source not found.** and Figure 3). The dry soil had lower Φ_P(II)_ and higher Φ_NPQ_ during the whole season based on polynomial regression (Figure 5). Water stress has been associated with sustained NPQ dissipation mechanism, with PSII core rearrangement or degradation (Demmig-Adams et al., 1996). Acclimation to sustained water stress conditions with significantly lower T_r_ and g_s_ (Table 2) can be identified by changes toward higher T_opt_ (O’sullivan et al., 2017; Yamori et al., 2014; Zheng et al., 2018) as it happened in our experiment where T_opt_ was also around 2°C to 3°C higher than in the wet scenario in all data subset selections. However, acclimation of photosynthetic enzymatic activity to higher temperatures has also been associated with reduced photosynthetic efficiency (Bagley et al., 2015) translating in lower yields (Luan et al., 2021) which could explain lower yields in our case due to acclimation to higher water stress, reducing g_s_ and T_r_, showing also slightly higher increases T_L_.

### Thermoregulation under heat stress days

We found two critical aspects reducing thermoregulation capacity in hot days that are different from the growing season thermoregulation: (i) the magnitude of the heat stress, T_shock_, that depresses maximum photochemical yield F_v_/F_m_ but inhibited only moderately Φ_P(II)_ (Figure 7) and (ii) limited soil water availability that reduces transpiration cooling via stomatal control. It can be expected that if the T_shock_ becomes large enough, those two effects will counteract the overall positive effect of T_L_ approaching T_opt_ that increases photochemical efficiency, identified by higher season averages of Φ_P(II)_ and lower Φ_NPQ_ in hot versus normal days (Table 2).

Plant responses to sudden increases in T_air_ differed from those related to gradual variation in T_air_ across the season. Our results for hot days were similar to Tsonev and Hikosaka (2003) with inhibition of F_v_/F_m_. Relative decreases in F_v_/F_m_ indicate photoinactivation of PSII. F_v_/F_m_ has been also described as a quick but unspecific stress indicator that has also been used to assess thermal stress (Geange et al., 2020). Indirect effects of heat could damage enzymes or membranes outside photosystem II, involving several processes (Murchie and Lawson, 2013; Tsonev and Hikosaka, 2003). In contrast to F_v_/F_m_, responses in Φ_P(II)_ involve just the biochemistry light reactions (e.g. carboxylation) (Baker, 2008). This could explain significantly higher Φ_P(II)_ comparing averages of hot days and normal days (Table 2), as plants reduce NPQ while enhance photosynthesis approaching the T_opt_ with warmer T_air_ (Tan et al., 2017; Zheng et al., 2018).

Note that our study aimed to investigate responses to absolute climatic extremes in T_air_ happening in the North China plain based on historical maxima of 15 years’ climatology in Beijing. We did not aim to find plant’s thermal limits, having maximum T_air_ in hot days set at 38°C (Figure 2). Most likely, due to the short duration and limited maximum T_air_ of the hot days in our experiment, the thermal safety margins were not passed. This is supported by observed significantly lower average values of Φ_NPQ_ during hot days compared to normal days (Table 2), contrary to a recent study, where during heat stress plants increased dissipation through NPQ causing also change in the allocation of energy towards SIF (Martini et al., 2022). During hot days, the rapid increase in T_air_ seems to affect photochemistry more than the absolute value of T_L_. When T_shock_ was large enough, flexible NPQ dissipation is activated, leading to potential declines in photosynthetic efficiency (Porcar-Castell et al., 2004). This is suggested in our dataset by reduced F_v_/F_m_ and moderate but significant increases in Φ_NPQ_ and decreases in Φ_p(II)_ (Figure 7) in agreement with the results of Martini et al., (2022). In our study, Φ_NPQ_ and Φ_p(II)_ for plants in hot days under drought conditions did not seem sensitive to changes in T_shock_ as they were already functioning with a lower photochemical efficiency from the sustained effect of photoinhibition described before with not clear responses in fluorescence (Figure 7).

Considering the range of temperatures of our study, our results fit with those of Crafts-Brander and Salvucci et al., (2002) who found that heat stress in maize reduced F_v_/F_m_, but this inhibition was marginal until T_L_ exceeded 42.5°C. Net photosynthesis was only inhibited at T_L_ above 38°C, and inhibition was much more severe when the temperature increased rapidly as in our hot days rather than gradually as during the season normal days where Rubisco activation could acclimate. It can be expected for temperatures way beyond T_opt_ in our experiment that plants will increase Φ_NPQ_ as indicated by the polynomial regression in Figure 5.

High evaporative cooling (expressed by EF being close to 1) was a key mechanism in hot days for thermoregulation but was lower for plants under soil drought as shown by hysteresis curves (Figure 8). Thus, when soil water supply was limited this capacity to cool was reduced due to a larger stomatal closure that would avoid compromising plant water status (Mathews and Lawson, 2019). This strategy to avoid catastrophic water loss, even though demands for photosynthesis should be still high, as the temperatures were not far from T_opt_, reflects the hierarchal response of one signal over-riding others (Moore et al., 2021).

In fact, stomatal regulation integrates the effect of changes in photosynthetic efficiency, water availability and atmospheric dryness (Tuzet et al., 2003). Several studies have suggested that g_s_ is driven by warmer T_air_. However, its response under heat stress is still uncertain. While some studies showed an increase of g_s_ with T_air_ increase, others found that stomata closed to warmer conditions or even had not response to T_air_ changes (see reviews in Urban et al., 2009; Moore et al., 2020). One explanation to these controversial responses is that we must consider not only the crop type and environmental conditions, but also the T_opt_ and how is T_L_ with respect to T_opt_. In our study, above T_opt_, we observed a significant decrease in g_s_ as water stress increased that should produce metabolic disruptions, and lower photosynthesis from restricted CO_2_ diffusion (Moore et al., 2021).

A key point is that despite the closure in g_s_, T_r_ or EF did not change as a season average (Table 2) or subdaily (Figure 8). This is because higher VPD due to high temperatures increases the leaf–atmosphere diffusion gradient, driving greater transpiration. The higher VPD should trigger stomatal closure to maintain plant water status protecting leaves from inhibitory short-term dehydration at the cost of small decreases in daily carbon gain (Field et al., 1982). Although generalizing stomatal response to changes in T_L_ is complicated, in our study when considering responses to VPD the responses seem unified (Figure 6) with a minimum of around 30 hPa and an optimum of around 14 hPa. Above VPD>35 hPa, g_s_ increases which can be considered an indicator of decoupling between g_s_ and photosynthesis (indicated in our case by inhibited F_v_/F_m_) as shown in other studies (Craft-Brander & Salvucci, 2002; Urban et al., 2017) whereby stomata open to increase leaf cooling despite the suppression of photosynthesis. This is contrary to the optimal stomatal hypothesis where stomata maximize photosynthesis while minimizing water losses (Moore et al., 2021).

## 5. Conclusion

This study showed that maize leaves thermoregulate under wet and dry soil scenarios as an adaptive mechanism to optimize photosynthesis around T_opt_. During the average growing season in controlled conditions, maize plants behaved as limited homeotherms, with T_eq_ being close to T_opt_, presenting a lower capacity to thermoregulate in the dry soil scenario, where a conservative water use reduced photosynthesis, g_s_ and T_r_ impacting final crop yields despite acclimation to sustained stress indicated by a higher T_opt_.

We propose for the first time a novel mechanism to explain thermoregulation based on the heat dissipation from photosynthesis or non-photochemical quenching (NPQ) that could open the path to estimate photosynthetic efficiency from thermal remote sensing in the future. NPQ dissipation acts as a negative feedback mechanism from photosynthesis supporting warming and leaf thermoregulation towards T_opt_ in suboptimal conditions. Close and above T_opt_ the main thermoregulation mechanism was stomatal regulation for transpiration cooling while NPQ was minimal.

Our results showed that plants coped better with heat if the T_air_ increase was not as sudden as in heatwaves, indicating that plants could slowly acclimate to rising temperatures as the season progresses. In the context of future climate in the North China Plain, our results suggest that even though a higher average T_air_ might bring maize photosynthesis closer to a thermal optimum, a higher intensity and/or frequency of extreme diurnal oscillations would reduce crop performance as shown by reductions in maximum photochemical yield, F_v_/F_m_, especially under water scarcity conditions.

## Author Contributions

Conceptualization: M.G., X.M., S.L; Methodology: M.G., X.M., V.S.-P, S.L., and L.H.; Measurements: V.S.-P, and L.H.; Investigation, data curation, formal analysis: V.S.-P., M.G., X.M., S.L., T.N.M. and H.J.; Resources: X.M., S.L., and M.G.; Supervision, M.G., X.M, S.L., and; T.N.M. Original draft, M.G. and V.S.-P. equal contribution; Writing—review, editing, all authors. All authors have read and agreed to the published version of the manuscript.

## Data availability

Data are available to researchers under a reasonable request from M.G or X.M.

## Acknowledgments

This article has benefited from the COST Action CA17134 “Optical synergies for spatiotemporal SENsing of Scalable ECOphysiological traits” (SENSECO. The authors would like to thank Pablo Miguel Martínez, Qiutan Chen, Enmin Liu, Lihu Yang and Xuejuan Chen for help on experimental set up and measurements.

## List of Supplementary data (Tables and Figures)

The following supplementary data are available at JXB online.

*Table* S1. Experiment management: irrigation and fertilization applied during the experiment in wet (100% Surface field capacity, SFC) and dry (40% SFC) lysimeters. DAS = days after sowing.

*Figure S1*. Estimates of daily mean evapotranspiration (ET, mmday^−1^) for wet (blue dots and continuous line) and dry (red starts and dashed line) soil lysimeters (scenarios). Irrigation amounts are indicated as well as the day that VPD changed. DAS = days after sowing. Vertical yellow lines indicate hot days or moderate heatwaves.

*Figure S2*. Air temperature (T_air_) vs leaf temperature from thermocouples (T_L_) for a) midday measurements between and for b) mean diurnal values between 6:00 a.m. and 7:00 p.m. of wet soil in blue (dots) and dry soil scenario in red (crosses). First order regressions are shown. The tables show regression parameters: slope (β), intercept, coefficient of determination (R^2^), p value (Wilcoxon test with Bonferroni correction, *p <= 0.05, **p <= 0.01, *** p <= 0.001 and ns = no significant), upper and lower 95% confidence interval (CI) values and the leaf-equivalence temperature (T_eq_), where T_air_=T_L_ based on regression. Heatwaves or hot days are yellow shaded.

*Figure S3*. Relationships among quantum yields. Quantum yields of photochemistry (Φ_P(II)_) vs. quantum yield of non-photochemical quenching (Φ_NPQ_) and with quantum yield of fluorescence (Φ_F_) in the secondary axis for a) wet soil moisture scenario and b) dry soil moisture scenario. c) and d) Relationship between Φ_F_ and Φ_P(II)_ for wet and dry soil scenarios, respectively. All figures show coefficient of determination (R^2^) and p-value of linear regression between quantum yields (Wilcoxon test with Bonferroni correction, *p <= 0.05, **p <= 0.01, *** p <= 0.001 and ns = no significant).

*Figure S4*. Subdaily averages of air temperature (T_air_), vapor pressure deficit (VPD), leaf temperature from thermocouples (T_L_), leaf temperature excess (ΔT), stomatal conductance (g_s_) and leaf transpiration rate (T_r_) across the season for normal days and hot days. Parameters in wet soil are given by a continuous blue line and in dry soil scenario with dashed red line. Numbers in the upper left corner show the % increase (↑) or decrease (↓) between normal days and heatwaves at 13.00 h.

## References

Aslam, M., Maqbool, M.A., Cengiz, R. 2015. Drought Stress in Maize (Zea mays L.) Effects, Resistance, Mechanisms, Global Achievements and Biological Strategies for Improvement. Cham: Springer. ISBN: 978-3-319-25440-1. https://doi.org/10.1007/978-3-319-25442-5

Baker, N.R. 2008. Chlorophyll Fluorescence: A Probe of Photosynthesis In Vivo. Annual Review of Plant Biology. 59, 89–113. https://doi.org/10.1146/annurev.arplant.59.032607.092759

Blonder B, Michaletz ST. 2018. A model for leaf temperature decoupling from air temperature. Agricultural and Forest Meteorology 262, 354–360.

Bonan, G. 2015. Ecological climatology: concepts and applications, Third Ed. ed. Cambridge University Press, Center for Atmospheric Research, Boulder, Colorado. https://doi.org/10.21425/f58433332

Brás, T.A., Seixas, J., Carvalhais, N., Jägermeyr, J. 2021. Severity of drought and heatwave crop losses tripled over the last five decades in Europe. Environmental Research Letters 16, 065012.

Buchner, O., Karadar, M., Bauer, I., Neuner, G., 2013. A novel system for in situ determination of heat tolerance of plants: First results on alpine dwarf shrubs. Plant Methods 9, 1. https://doi.org/10.1186/1746-4811-9-7

Campbell, G.S., Norman, J.M., 1998. An Introduction to Environmental Biophysics, 2nd ed, Springer. https://doi.org/10.2134/jeq1977.00472425000600040036x

Chen, X., Mo, X., Hu, S., Liu, S. 2019. Relationship between fluorescence yield and photochemical yield under water stress and intermediate light conditions. Journal of Experimental Botany 70, 301–313. https://doi.org/10.1093/jxb/ery341

Crafts-Brandner, S.J., Salvucci, M.E. 2002. Sensitivity of Photosynthesis in a C4 Plant, Maize, to Heat Stress. Plant Physiology 129, 1773–1780. https://doi.org/10.1104/pp.002170.or

Das, A., Eldakak, M., Paudel, B., Kim, D., Hemmati, H., Basu, C., Rohila, J.S., 2016. Leaf Proteome Analysis Reveals Prospective Drought and Heat Stress Response Mechanisms in Soybean. BioMed Research International 2016, 23. https://doi.org/10.1155/2016/6021047

De Boeck, H.J., Dreesen, F.E., Janssens, I.A., Nijs, I. 2010. Climatic characteristics of heat waves and their simulation in plant experiments. Global Change Biology 16, 1992–2000. https://doi.org/10.1111/j.1365-2486.2009.02049.x

Demmig-Adams B, Adams III WW. 2006. Photoprotection in an ecological context: the remarkable complexity of thermal energy dissipation. New Phytologistogist 172, 11–21.

Drake, J.E., Harwood, R., Vårhammar, A., Barbour, M.M., Reich, P.B., Barton, C.V.M., Tjoelker, M.G. 2020. No evidence of homeostatic regulation of leaf temperature in Eucalyptus parramattensis trees: integration of CO2 flux and oxygen isotope methodologies. New Phytologist 1511–1523. https://doi.org/10.1111/nph.16733

Drake, J.E., Tjoelker, M.G., Vårhammar, A., Medlyn, B.E., Reich, P.B., Leigh, A., Pfautsch, S., Blackman, C.J., López, R., Aspinwall, M.J., Crous, K.Y., Duursma, R.A., Kumarathunge, D., De Kauwe, M.G., Jiang, M., Nicotra, A.B., Tissue, D.T., Choat, B., Atkin, O.K., Barton, C.V.M., 2018. Trees tolerate an extreme heatwave via sustained transpirational cooling and increased leaf thermal tolerance. Global Change Biology 24, 2390–2402. https://doi.org/10.1111/gcb.14037

Duursma, R.A., Barton, C.V.M., Lin, Y.S., Medlyn, B.E., Eamus, D., Tissue, D.T., Ellsworth, D.S., McMurtrie, R.E. 2014. The peaked response of transpiration rate to vapour pressure deficit in field conditions can be explained by the temperature optimum of photosynthesis. Agricultural and Forest Meteorology 189-190, 2–10. https://doi.org/10.1016/j.agrformet.2013.12.007

Endo, T., Uebayashi, N., Ishida, S., Ikeuchi, M., & Sato, F. 2014. Light energy allocation at PSII under field light conditions: how much energy is lost in NPQ-associated dissipation?. Plant Physiology and Biochemistry 81, 115–120.

Farella MM, Fisher JB, Jiao W, Key KB, Barnes ML. 2022. Thermal remote sensing for plant ecology from leaf to globe. Journal of Ecology 110, 1996–2014.

EURAMET, 2011. Calibration of Thermocouples. Calibration Guide. Eurpoean Association of National Metrology Institutes.

FAO, 2020. Food and Agriculture Organization of the Unitated States. Land & Water: Crop Water Information. [WWW Document]. URL http://www.fao.org/land-water/databases-and-software/crop-information/en/ (accessed 6.17.20).

Fauset, S., Freitas, H.C., Galbraith, D.R., Sullivan, M.J.P., Aidar, M.P.M., Joly, C.A., Phillips, O.L., Vieira, S.A., Gloor, M.U. 2018. Differences in leaf thermoregulation and water use strategies between three co-occurring Atlantic forest tree species. Plant, Cell and Environment 41, 1618–1631. https://doi.org/10.1111/pce.13208

Geange, S.R., Arnold, P.A., Catling, A.A., Coast, O., Cook, A.M., Gowland, K.M., Leigh, A., Notarnicola, R.F., Posch, B.C., Venn, S.E., Zhu, L., Nicotra, A.B. 2020. The thermal tolerance of photosynthetic tissues: a global systematic review and agenda for future research. New Phytologist 229, 2497–2513. https://doi.org/10.1111/nph.17052

Gerhards, M., Rock, G., Schlerf, M., Udelhoven, T. 2016. Water stress detection in potato plants using leaf temperature, emissivity, and reflectance. International Journal of Applied Earth Observation and Geoinformation 53, 27–39. https://doi.org/10.1016/j.jag.2016.08.004

Grossiord, C., Buckley, T.N., Cernusak, L.A., Novick, K.A., Poulter, B., Siegwolf, R.T.W., Sperry, J.S., McDowell, N.G. 2020. Plant responses to rising vapor pressure deficit. New Phytologist 226, 1550–1566. https://doi.org/10.1111/nph.16485

Guha, A., Han, J., Cummings, C., Mclennan, D.A., Warren, J.M. 2018. Differential ecophysiological responses and resilience to heat wave events in four co-occurring temperate tree species. Environmental Research Letters 13, 065008.

Hatfield, J.L., Boote, K.J., Kimball, B.A., Ziska, L.H., Izaurralde, R.C., Ort, D., Thomson, A.M., Wolfe, D. 2011. Climate impacts on agriculture: Implications for crop production. Agronomy Journal 103, 351–370.

Hirotsu, N., Makino, A., Ushio, A., Mae, T. 2004. Changes in the Thermal Dissipation and the Electron Flow in the Water – Water Cycle in Rice Grown Under Conditions of Physiologically Low Temperature. Plant Cell Physiology 45, 635–644.

Humplík, J.F., Lazár, D., Husičková, A., Spíchal, L. 2015. Automated phenotyping of plant shoots using imaging methods for analysis of plant stress responses - A review. Plant Methods 11, 1–10. https://doi.org/10.1186/s13007-015-0072-8

IPCC, 2020. Summary for Policymakers. In: Climate Change and Land: an IPCC special report on climate change, desertification, land degradation, sustainable land management, food security, and greenhouse gas fluxes in terrestrial ecosystems. https://doi.org/10.1002/9781118786352.wbieg0538

Jagadish SVK, Way DA, Sharkey TD. 2021. Plant heat stress: concepts directing future research. Plant, Cell & Environment.

Jimenez, I.M., Larkum, A.W.D., Ralph, P.J., Kühl, M. 2012. Thermal effects of tissue optics in symbiont-bearing reef-building corals. Limnology and Oceanography 57, 1816–1825. https://doi.org/10.4319/lo.2012.57.6.1816

Jones, H.G. and Rotenberg, E. 2011. Energy, Radiation and Temperature Regulation in Plants. In eLS, John Wiley & Sons, Ltd (Ed.). https://doi.org/10.1002/9780470015902.a0003199.pub2

Kaňa, R., Vass, I. 2008. Thermoimaging as a tool for studying light-induced heating of leaves. Correlation of heat dissipation with the efficiency of photosystem II photochemistry and non-photochemical quenching. Environmental and Experimental Botany 64, 90–96. https://doi.org/10.1016/j.envexpbot.2008.02.006

Killi, D., Bussotti, F., Raschi, A., Haworth, M. 2017. Adaptation to high temperature mitigates the impact of water deficit during combined heat and drought stress in C3 sunflower and C4 maize varieties with contrasting drought tolerance. Physiologia Plantarum 159, 130–147. https://doi.org/10.1111/ppl.12490

Lambers, H., Chapin III, F.S., Pons, T.L. 2008. Plant Physiologyogical Ecology, Second Ed. ed. Springer Science + Bussiness Media B.V. https://doi.org/10.1007/978-0-387-78341-3

Lee, J., Berry, J.A., van der Tol, C., Yang, X.I., Guanter, L., Damm, A., Baker, I., Frankenberg, C. 2015. Simulations of chlorophyll fluorescence incorporated into the Community Land Model version 4. Global Change Biology. 21, 3469–3477. https://doi.org/10.1111/gcb.12948

Leinonen, I., Jones, H.G. 2004. Combining thermal and visible imagery for estimating canopy temperature and identifying plant stress. Journal of Experimental Botany 55, 1423–1431. https://doi.org/10.1093/jxb/erh146

Li, Y., Guan, K., Peng, B., Franz, T.E., Wardlow, B., Pan, M. 2020. Quantifying irrigation cooling benefits to maize yield in the US Midwest. Global Change Biology 26: 3065–3078. https://doi.org/10.1111/gcb.15002

Liu, S., Mo, X., Lin, Z., Xu, Y., Ji, J., Wen, G., Richey, J. 2010. Crop yield responses to climate change in the Huang-Huai-Hai Plain of China. Agricultural Water Management 97, 1195–1209. https://doi.org/10.1016/j.agwat.2010.03.001

Lobell, D.B., Hammer, G.L., McLean, G., Messina, C., Roberts, M.J., Schlenker, W. 2013. The critical role of extreme heat for maize production in the United States. Nature Climate Change 3, 497–501. https://doi.org/10.1038/nclimate1832

Luan, X., Bommarco, R., Scaini, A., Vico, G. 2021. Combined heat and drought suppress rainfed maize and soybean yields and modify irrigation benefits in the USA. Environmental Research Letters 16, 064023. https://doi.org/10.1088/1748-9326/abfc76

Luan, X., Vico, G. 2020. Canopy temperature and heat stress are increased by compound high air temperature and water stress, and reduced by irrigation - A modeling analysis. Hydrology and Earth Systems Sciences 25, 1411–1423. https://doi.org/10.5194/hess-25-1411-2021 https://doi.org/10.5194/hess-2020-549

Mahan JR, Upchurch DR. 1988. Maintenance of constant leaf temperature by plants—I. Hypothesis- limited homeothermy. Environmental and Experimental Botany 28, 351–357.

Martini, D., Sakowska, K., Wohlfahrt, G., Pacheco-labrador, J., Tol, C. Van Der, Porcar-Castell, A., Magney, T.S., Carrara, A., Colombo, R., El-Madany, T.S., Gonzalez-Cascon, R., Martin, M.P., Julitta, T., Moreno, G., Rascher, U., Reichstein, M., Rossini, M., Migliavacca, M. 2022. Heatwave breaks down the linearity between sun-induced fluorescence and gross primary production. New Phytologist. 233, 2415–2428. https://doi.org/10.1111/nph.17920

Michaletz, S.T., Weiser, M.D., McDowell, N.G., Zhou, J., Kaspari, M., Helliker, B.R., Enquist, B.J., 2016. The energetic and carbon economic origins of leaf thermoregulation. Nature Plants 16129. https://doi.org/10.1038. https://doi.org/10.1038/nplants.2016.129

Michaletz, S.T., Weiser, M.D., Zhou, J., Kaspari, M., Helliker, B.R., Enquist, B.J. 2015. Plant Thermoregulation: Energetics, Trait-Environment Interactions, and Carbon Economics. Trends in Ecology and Evolution 30, 714–724. https://doi.org/10.1016/j.tree.2015.09.006

Mishra, V., Cherkauer, K.A. 2010. Retrospective droughts in the crop growing season : Implications to corn and soybean yield in the Midwestern United States. Agricultural and Forest Meteorology 150, 1030–1045. https://doi.org/10.1016/j.agrformet.2010.04.002

Moore, C.E., Meacham-Hensold, K., Lemonnier, P., Slattery, R.A., Benjamin, C., Bernacchi, C.J., Lawson, T., Cavanagh, A.P. 2021. The effect of increasing temperature on crop photosynthesis: from enzymes to ecosystems. Journal of Experimental Botany 72, 2822–2844. https://doi.org/10.1093/jxb/erab090

Murchie, E.H., Lawson, T. 2013. Chlorophyll fluorescence analysis: A guide to good practice and understanding some new applications. Journal of Experimental Botany 64, 3983–3998. https://doi.org/10.1093/jxb/ert208

O’Sullivan, O.S., Heskel, M.A., Reich, P.B., Tjoelker, M.G., Weerasinghe, L.K., Penillard, A., Zhu, L., Egerton, J.J.G., Bloomfield, K.J., Creek, D., Bahar, N.H.A., Griffin, K.L., Hurry, V., Meir, P., Turnbull, M.H., Atkin, O.K. 2017. Thermal limits of leaf metabolism across biomes. Global Change Biology 23, 209–223. https://doi.org/10.1111/gcb.13477

Porcar-Castell A, Tyystjärvi E, Atherton J, Van Der Tol C, Flexas J, Pfündel EE, Moreno J, Frankenberg C, Berry JA. 2014. Linking chlorophyll a fluorescence to photosynthesis for remote sensing applications: mechanisms and challenges. Journal of Experimental Botany 65, 4065–4095.

Roy, J., Rineau, F., De Boeck, H.J., Nijs, I., Pütz, T., Abiven, S., Arnone, J.A., Barton, C.V.M., Beenaerts, N., Brüggemann, N., Dainese, M., Domisch, T., Eisenhauer, N., Garré, S., Gebler, A., Ghirardo, A., Jasoni, R.L., Kowalchuk, G., Landais, D., Larsen, S.H., Leemans, V., Le Galliard, J.F., Longdoz, B., Massol, F., Mikkelsen, T.N., Niedrist, G., Piel, C., Ravel, O., Sauze, J., Schmidt, A., Schnitzler, J.P., Teixeira, L.H., Tjoelker, M.G., Weisser, W.W., Winkler, B., Milcu, A. 2021. Ecotrons: Powerful and versatile ecosystem analysers for ecology, agronomy and environmental science. Global Change Biology 27, 1387–1407. https://doi.org/10.1111/gcb.15471

Ruiz-Vera, U.M., Siebers, M.H., Drag, D.W., Ort, D.R., Bernacchi, C.J. 2015. Canopy warming caused photosynthetic acclimation and reduced seed yield in maize grown at ambient and elevated [CO2]. Global Change Biology 21, 4237–4249. https://doi.org/10.1111/gcb.13013

Schulze, ED., Beck, E., Buchmann, N., Clemens, S., Müller-Hohenstein, K., Scherer-Lorenzen, M. 2019. Thermal Balance of Plants and Plant Communities. In: Plant Ecology. Springer, Berlin, Heidelberg. https://doi.org/10.1007/978-3-662-56233-8_9

Seneviratne, S.I., Nicholls, N., Easterling, D., Goodess, C.M., Kanae, S., Kossin, J., Luo, Y., Marengo, J., Mc Innes, K., Rahimi, M., Reichstein, M., Sorteberg, A., Vera, C., Zhang, X., Rusticucci, M., Semenov, V., Alexander, L. V., Allen, S., Benito, G., Cavazos, T., Clague, J., Conway, D., Della-Marta, P.M., Gerber, M., Gong, S., Goswami, B.N., Hemer, M., Huggel, C., Van den Hurk, B., Kharin, V. V., Kitoh, A., Klein Tank, A.M.G., Li, G., Mason, S., Mc Guire, W., Van Oldenborgh, G.J., Orlowsky, B., Smith, S., Thiaw, W., Velegrakis, A., Yiou, P., Zhang, T., Zhou, T., Zwiers, F.W. 2012. Changes in climate extremes and their impacts on the natural physical environment. In: Managing the Risks of Extreme Events and Disasters to Advance Climate Change Adaptation [Field, C.B., V. Barros, T.F. Stocker, D. Qin, D.J. Dokken, K.L. Ebi, M.D. Mastrandrea, K.J. Mach, G.-K. Plattner, S.K. Allen, M. Tignor, and P.M. Midgley (eds.)]. A Special Report of Working Groups I and II of the Intergovernmental Panel on Climate Change (IPCC). Cambridge University Press, Cambridge, UK, and New York, NY, USA, pp. 109–230

Sobejano-Paz, V., Mikkelsen, T.N., Baum, A., Mo, X., Liu, S., Köppl, C.J., Johnson, M.S., Gulyas, L., García, M. 2020. Hyperspectral and Thermal Sensing of Stomatal Conductance, Transpiration, and Photosynthesis for Soybean and Maize under Drought. Remote Sensing 12, 3182. https://doi.org/10.3390/rs12193182

Still, C., Powell, R., Aubrecht, D., Kim, Y., Helliker, B., Roberts, D., Richardson, A.D., Goulden, M. 2019. Thermal imaging in plant and ecosystem ecology: applications and challenges. Ecosphere 10(6): e02768.https://doi.org/10.1002/ecs2.2768

Still, C. J., Page, G., Rastogi, B., Griffith, D. M., Aubrecht, D. M., Kim, Y., … & Richardson, A. D. 2022. No evidence of canopy-scale leaf thermoregulation to cool leaves below air temperature across a range of forest ecosystems. Proceedings of the National Academy of Sciences, 119(38), e2205682119.

Tan, L., Liu, S., Mo, X., Lin, Z., Hu, S., Qiao, Y., Liu, E., Song, X. 2017. Responses of maize ecosystem to warming based on WATDPED experiment under near-field condition. Ecological Indicators 81, 390–400.

Teuling, A.J, 2018. A hot future for European droughts. Nature Climate Change 8, 364–365. https://doi.org/10.1038/s41558-018-0154-5

Tiwari, Y.K., Yadav, S.K. 2019. High Temperature Stress Tolerance in Maize (Zea mays L.): Physiological and Molecular Mechanisms. Journal of Plant Biology 62, 93–102. https://doi.org/10.1007/s12374-018-0350-x

Tsonev, T.D., Hikosaka, K. 2003. Contribution of Photosynthetic Electron Transport, Heat Dissipation, and Recovery of Photoinactivated Photosystem II to Photoprotection at Different Temperatures in Chenopodium album Leaves 44, 828–835.

Tuzet, A., Perrier, A., Leuning, R. 2003. A coupled model of stomatal conductance, photosynthesis and transpiration. Plant, Cell and Environment 26, 1097–1116. https://doi.org/10.1007/s11099-008-0110-0

Urban, J., Ingwers, M.W., McGuire, M.A., Teskey, R.O., 2017. Increase in leaf temperature opens stomata and decouples net photosynthesis from stomatal conductance in Pinus taeda and Populus deltoides x nigra. Journal of Experimental Botany 68, 1757–1767. https://doi.org/10.1093/jxb/erx052

Vallat, R. 2018. Pingouin: statistics in Python. J. Open Source Software 3, 1026. https://doi.org/10.21105/joss.01026

Van der Tol, C., Berry, J.A., Campbell, P.K.E., Rascher, U. 2014. Models of fluorescence and photosynthesis for interpreting measurements of solar-induced chlorophyll fluorescence. Journal of Geophysical Research, Biogeosciences 119, 2312–2327. https://doi.org/10.1002/2014JG002713.We

Vico, G., Way, D.A., Hurry, V., Manzoni, S. 2019. Can leaf net photosynthesis acclimate to rising and more variable temperatures? Plant, Cell and Environment 42, 1913–1928. https://doi.org/10.1111/pce.13525

Wu, Y., Song, X., Ma, Y., Bu, H., Yang, S., Han, D., Zhang, Y., 2017. Multi-temporal variation in water consumption of summer maize as determined by the Water Transformation Dynamical Processes Experimental Device Hydrology Research 48. https://doi.org/10.2166/nh.2016.021

Yamori, W., Hikosaka, K., Way, D.A., 2014. Temperature response of photosynthesis in C3, C4, and CAM plants : temperature acclimation and temperature adaptation. Photosynthis Research 101–117. https://doi.org/10.1007/s11120-013-9874-6

Zheng, Y.P., Li, R.Q., Guo, L.L., Hao, L.H., Zhou, H.R., Li, F., Peng, Z.P., Cheng, D.J., Xu, M. 2018. Temperature Responses of Photosynthesis and Respiration of Maize (Zea mays) Plants to Experimental Warming. Russian Journal of Plant Physiology 65, 524–531. https://doi.org/10.1134/S1021443718040192

